# Adaptation and the Parliament of Genes

**DOI:** 10.1101/526970

**Authors:** Thomas W. Scott, Stuart A. West

## Abstract

Fields such as behavioural and evolutionary ecology are built on the assumption that natural selection leads to organisms that behave as if they are trying to maximise their fitness. However, there is considerable evidence for selfish genetic elements that change the behaviour of individuals to increase their own transmission. How can we reconcile these contradictions? We found theoretically that, when selfish genetic elements have a greater impact at the individual level, they are more likely to be suppressed, and suppression spreads more quickly. Consequently, selfish genetic elements will either have a minor impact at the individual level, or tend to be suppressed. In addition, we found that selection on selfish genetic elements favours higher levels of distortion. Consequently, selfish genetic elements will tend to evolve to make themselves more likely to be suppressed. Overall, our results suggest that even when there is the potential for considerable genetic conflict, this will often have negligible impact at the individual level.

## Introduction

There is a clear contradiction between major branches of modern evolutionary biology. On the one hand, fields such as behavioural and evolutionary ecology are based on the assumption that organisms will behave as if they are trying to maximise their fitness^1–4^. Models based on fitness maximisation are used to make predictions about the selective forces (reasons) for adaptation, and these are then tested empirically^5,6^. This approach has been phenomenally successful, explaining many aspects of behaviour, life history and morphology. For example, research on: foraging, resource competition, sexual selection, parental care, sex allocation, signalling and cooperation^7–12^.

However, on the other hand, there is considerable evidence for selfish genetic elements, which increase their own contribution to future generations at the expense of other genes in the same organism^13–16^. These ‘selfish genes’ distort traits away from the values that would maximise individual fitness, to increase their own transmission^14,17–21^. Evidence for such genetic conflict has been found across the tree of life, from simple prokaryotes to complex animals. The contradiction is that selfish genetic elements mess up individual fitness maximisation, and appear to be common. But individual fitness maximisation still appears to occur^22,23^. For example, research on sex allocation has provided phenomenal support for both the individual fitness maximisation approach, and the existence of selfish sex ratio distorters^9,14,24–26^.How can we resolve this contradiction?

Leigh^27^ provided a potential solution to this contradiction by suggesting that selfish genetic elements would be suppressed by the ‘parliament of genes’. Leigh’s argument was that, because selfish genetic elements reduce the fitness of the other genes in the organism, the rest of the genome will have a united interest in suppressing selfish genetic elements. Furthermore, because those other genes are far more numerous, they will be likely to win the conflict. Consequently, even when there is considerable potential for conflict within individuals, we would still expect fitness maximisation at the individual level^28–32^. Leigh^27^ demonstrated the plausibility of his argument by showing theoretically how a suppressor of a meiotic drive gene could be favoured.

However, there are a number of potential problems with the parliament of genes hypothesis. First, even if a parliament of genes could easily generate suppressors, whether a suppressor spreads can depend upon biological details such as any cost associated with the suppressor, the extent to which a selfish genetic element is distorting a trait, and the population frequency of that selfish genetic element^14,33–38^. Second, even if suppressors can spread, prolonged non-equilibrium trait distortion is possible if the spread of suppressors through populations is slow. Third, selfish genetic elements are themselves also under evolutionary pressure to reach a level of distortion that would maximise their transmission to the next generation – how will this influence the likelihood that they are suppressed?^31^ Fourth, segregation at suppressor loci might expose previously suppressed selfish genetic elements^34^.

We investigated the parliament of genes hypothesis theoretically. Our aim was to investigate the extent to which genetic conflict distorts traits away from the value that would maximise individual fitness. We found that: (i) the greater the level of distortion caused by a selfish genetic element, the more likely and the quicker it will be suppressed; (ii) selection on selfish genetic elements leads towards greater distortion, making them more likely to be suppressed. We found the same patterns with an illustrative model, and when examining three specific examples: selfish distortion of the sex ratio by an X chromosome driver; an altruistic helping behaviour encoded by an imprinted gene; and production of a cooperative public good encoded on a horizontally transmitted bacterial plasmid. Furthermore, we found close agreement when analysing scenarios with population genetic analyses and individual based simulations. Our results suggest that even when there is potential for considerable genetic conflict, it will have relatively little impact on traits at the individual level.

## Results

A selfish genetic element may be able to gain a propagation advantage through trait distortion (‘*distorter*’). Any part of the genome that does not gain the propagation advantage from the trait distortion will be selected to suppress the distorter. This collection of genes will comprise most of the genome, and so will constitute the majority within the parliament of genes^20^. We take into account the large size of this collection of genes by assuming that it is highly likely that a potential suppressor of a distorter can arise by mutation (high mutational accessibility). Consequently, we focus our analyses on when a distorter and its suppressor can spread. Prolonged trait distortion may be possible if a suppressor is hard to evolve, but our assumption of suppressor availability allows us to focus on the evolutionary dynamics that succeed the acquisition of a suppressor by mutation^31^.

Our overall aim is to assess, given the potential for suppression, the extent to which a distorter can distort the organism trait away from the optimum for individuals (the largest “coreplicon”^21^). In order to elucidate the selective forces in operation, we ask four questions in a step-wise manner, with increasing complexity:

1. In the absence of a suppressor, when can a distorter invade?
2. When can a costly suppressor of the distorter invade?
3. What are the overall consequences of distorter-suppressor dynamics for trait values, at the individual and population level, at evolutionary equilibrium and before equilibrium has been reached?
4. If the extent to which the distorter manipulates the organism trait can evolve, how will this influence the likelihood that it is suppressed, and hence the individual and population trait values?

### Illustrative Model

We assume an arbitrary trait that influences organism fitness. In the absence of distorters, all individuals have the trait value that maximises their individual fitness. The distorter manipulates the trait away from the individual optimum, to increase their own transmission to offspring. We assume a large population of diploid, randomly mating individuals. The aim of this model is to establish key aspects of the population genetics governing distorters and their suppressors, in an abstract setting. We will subsequently address analogous issues in three specific biological scenarios.

### (1) Spread of a Distorter

We consider a distorter, which we denote by *y*_*1*_, that is dominant and distorts an organism trait value by some positive amount, denoted by *k* (*k*>0). This distortion increases the transmission of the distorter to offspring. Specifically, the distorter (*y*_1_) drives at meiosis, in heterozygotes, against a non-distorter (*y*_*0*_), being passed into the proportion (1+*t*(*k*))/2 of offspring. *t*(*k*) denotes the transmission bias (0≤t(*k*)≤1) and is a monotonically increasing function of trait distortion 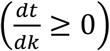.

Trait distortion leads to a fitness (viability) cost (*c*_*drive*_(*k*)) at the individual level, reducing an individual’s number of offspring from 1 to 1-*c*_*drive*_(*k*) (0≤*c*_*drive*_(*k*)≤1). Owing to distorter dominance, the fitness cost of trait distortion is borne by heterozygous as well as distorter-homozygous individuals. The fitness cost is a monotonically increasing function of trait distortion 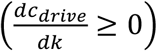. We assume that *t*(*k*) and *c*_*drive*_(*k*) do not change with population allele frequencies, but relax this assumption in our specific models.

We first ask what frequency the distorter will reach in the population in the absence of suppression. If we take *p* and *p’* as the population frequency of the distorter in two consecutive generations, then the population frequency of the distorter in the latter generation is:

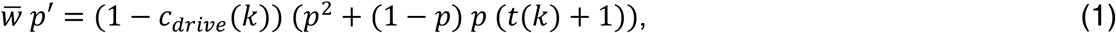

where 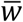 is the average fitness of individuals in the population in the current generation, and can be written in full as: 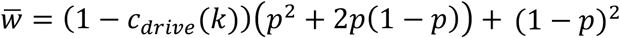. In Appendix 1 we show, with a population genetic analysis of equation (1),that the distorter will spread from rarity and reach fixation when *c*_*drive*_(*k*)<t(*k*)(1*-c*_*drive*_(*k*)). This shows that distortion will evolve when the number of offspring that the distorter gains as a result of distortion (t(*k*)(1*-c*_*drive*_(*k*))) is greater that the number of offspring bearing the distorter that are lost as a result of reduced individual fitness (*c*_*drive*_(*k*)).

### (2) Spread of an autosomal suppressor

We assume that the distorter (*y*_*1*_) can be suppressed by an unlinked autosomal allele (suppressor), denoted by *sup*. This suppressor (*sup*) is dominant and only expressed in the presence of the distorter (facultative), and its expression may lead to a fitness cost to the individual, *c*_*sup*_ (0≤*c*_*sup*_≤1)^39,40^. This cost can arise for multiple reasons, including energy expenditure, or errors relating to the use of gene silencing machinery, and is likely to be relatively low^41^. Gene silencing generally precedes the translation of the targeted gene, and so we assume that the costs of suppression (*c*_*sup*_) is independent of the amount of distortion caused by the distorter (*k*).

We can write recursions detailing the generational change in the frequencies of the four possible gametes, *y_0_*/+, *y_0_*/*sup*, *y*_*1*_/+, *y*_*1*_/*sup*, with the respective frequencies in the current generation denoted by *x*_*1*_, *x*_*2*_, *x*_*3*_ and *x*_*4*_, and the frequencies in the subsequent generation denoted by an appended dash (’):

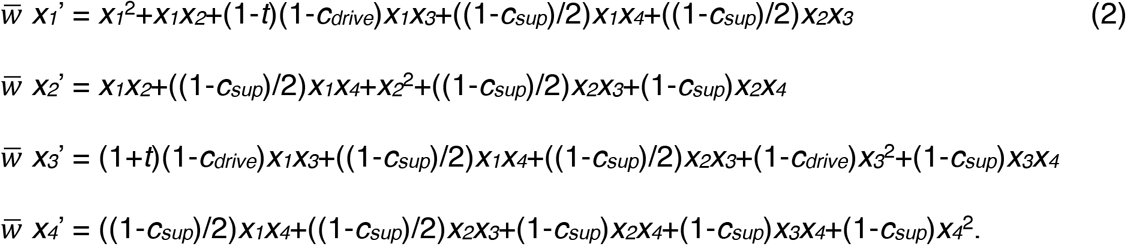

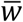 is the average fitness of individuals in the current generation, and equals the sum of the equations’ right-hand sides. In Appendix 2, we show, with a population genetic analysis of our system of equations (2), that a suppressor will spread from rarity if distortion, *k*, is greater than some threshold value at which the cost of suppression (*c*_*sup*_) is less than the cost of being subjected to trait distortion, *c*_sup_<c_drive_(*k*). A threshold with respect to the level of distortion (*k*) arises because the cost of trait distortion (*c*_*drive*_(*k*)) increases with greater distortion, but the cost of suppression (*c*_*sup*_) is constant.

### (3) Consequences for organism trait values

The extent of trait distortion at the individual level shows a discontinuous relationship with the strength of the distorter. When distortion is low, a suppressor will not spread (*c_sup_>c*_*drive*_(*k*)) and so the level of trait distortion at the individual level will increase with the level of trait distortion induced by the distorter (*k*). However, once a threshold is reached (*c_sup_<c*_*drive*_(*k*)), the suppressor spreads. We show in Appendix 3 that the spread of the suppressor causes the distorter (*y*_*1*_) to lose its selective advantage and be eliminated from the population, leading to an absence of distortion at the individual level.

Overall, these results suggest that, given a relatively low cost of suppression (*c*_*sup*_), the level of distortion observed at the individual level will either be low or absent. When a distorter is weak (low *k*), it will not be suppressed, but it will only have a small influence at the level of the individual. When a distorter is strong (high *k*), it will be suppressed and so there will be no influence at the level of the individual (Fig. 1a).

**Figure 1.**
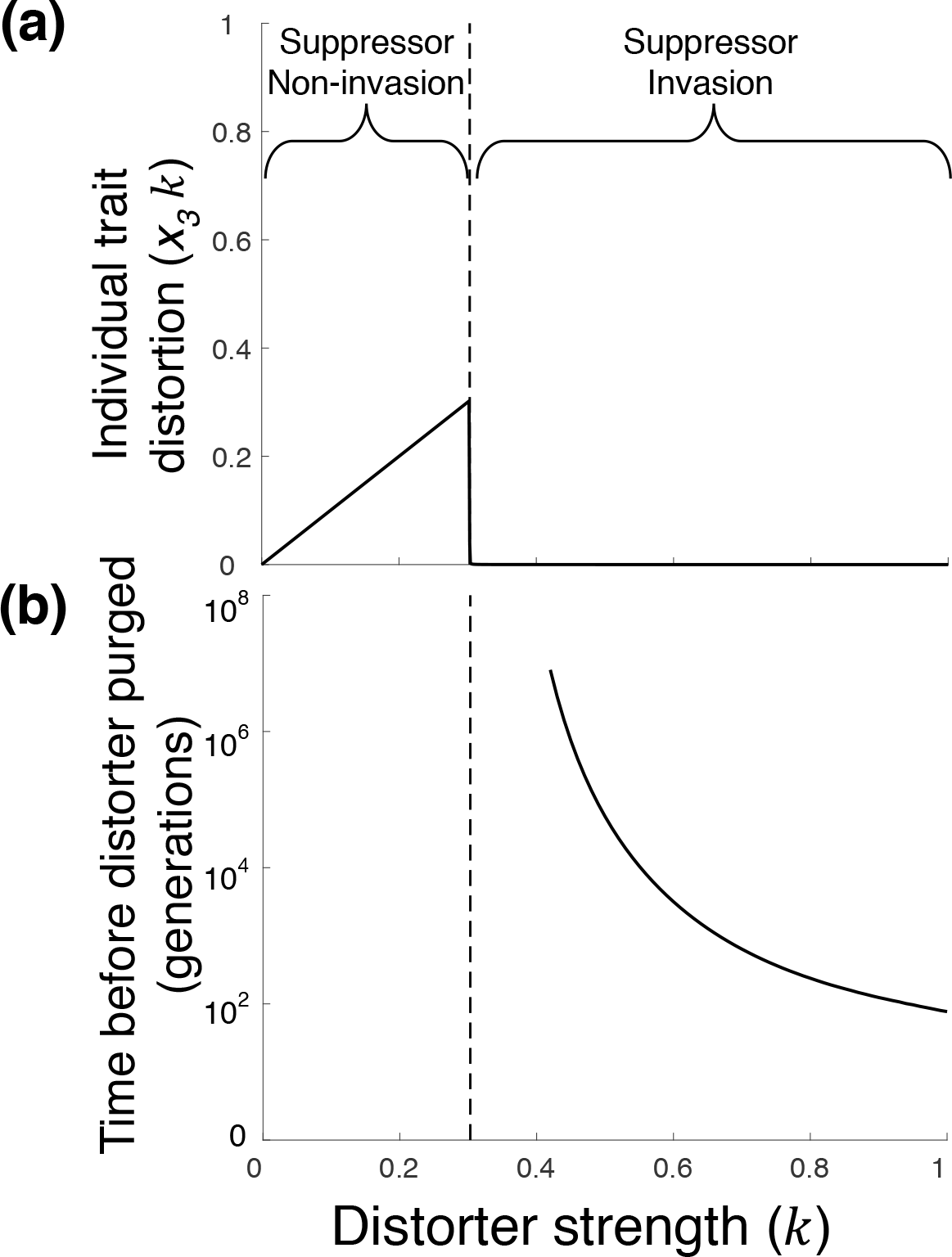
Distorter-suppressor dynamics and consequences for the organism. The trait distorter (*y*_*1*_) and its suppressor (*sup*) are introduced from rarity. In part (a), the resulting average trait distortion (*x*_3_*k*) is plotted at equilibrium, as the solid line, against the extent to which the distorter causes trait values to deviate from the individual optimum (*k*). Below a certain threshold strength (*c*_*sup*_ >*c*_*drive*_(*k*)), to the left of the dashed line, the suppressor does not invade, and so the resulting trait distortion increases with the strength of the distorter (*k*). However, above this threshold, the suppressor invades, and the distorter is purged, restoring the trait to the individual optimum. In part (b), the number of generations for distorters that are purged at equilibrium (having been suppressed), which lie to the right of the dashed line, is plotted on a log_10_ scale. Stronger distorters are purged more quickly than weaker distorters. The numerical solutions displayed graphically assume that the cost of suppression, the transmission benefit of distortion, and the individual fitness cost of distortion, are respectively given by: *c*_*drive*_ =0.15; *t*=0.87*k* and *c*_*drive*_ =0.9*k*^1.5^.

We also found that stronger distorters are suppressed more quickly (Fig. 1b). In Appendix 4, we numerically iterated our recursions to determine how many generations it takes for suppressors to reach equilibrium. As long as trait distortion continues to reduce individual fitness non-negligibly after suppression is favoured (such that 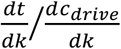 is not excessively high after *c_sup_<c*_*drive*_(*k*)), stronger distorters (higher *k*) are suppressed and purged more rapidly than weaker distorters, limiting the potential for non-equilibrium trait distortion (Fig. 1b).

### (4) Evolution of trait distortion

We then considered the consequence of allowing the level of trait distortion (*k*) to evolve. We assume a distorter (*y_1_*) that distorts by *k*, and then introduce a rare mutant (*y_2_*) that distorts by a different amount 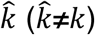. This mutant (*y_2_*) is propagated into the proportion 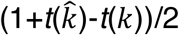 of the offspring of *y*_2_*y*_1_ heterozygotes, and into the proportion 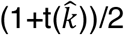 of the offspring of *y*_2_*y*_0_ heterozygotes. We assume that the stronger of the two distorters is dominant, but found similar results when assuming additivity (Appendix 5). We assume that the similarity in coding sequence and regulatory control means that the original distorter and the mutant are both suppressed by the same suppressor allele, at the same cost (*c_sup_*)49. In Appendix 5, we write the recursions that detail the generational frequency changes in the different possible gametes (*y*_*0*_/+, *y*_*0*_/*sup*, *y*_*1*_/+, *y*_*1*_/*sup*, *y*_*2*_/+, *y*_*2*_/*sup*).

We found that stronger mutant distorters 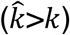 will invade from rarity when the marginal increase in offspring they are propagated into exceeds the marginal increase in offspring they are lost from as a result of reduced fitness 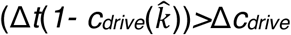, where Δ denotes marginal change 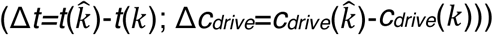. Consequently, if distortion is initially low, and successive mutant distorters are introduced, each deviating only slightly from the distorters from which they are derived (“δ-weak selection”^42^), invading distorters will approach a ‘target’ strength, denoted by *k*_*target*_. The target strength is that at which the marginal benefit of transmission is exactly counterbalanced by the marginal individual cost of reduced offspring, which occurs when 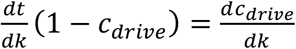. The target strength of distortion (*k*_*target*_) will therefore be greater if increased trait distortion (*k*) leads to a low rate of decrease in marginal transmission benefit 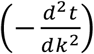 relative to the rate of increase in marginal individual cost 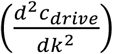 (Fig. 2b). If mutations are larger (strong selection), invading distorters may overshoot the target strength of distortion 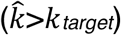. Weaker mutant distorters 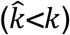 are recessive so cannot invade from rarity.

**Figure 2.**
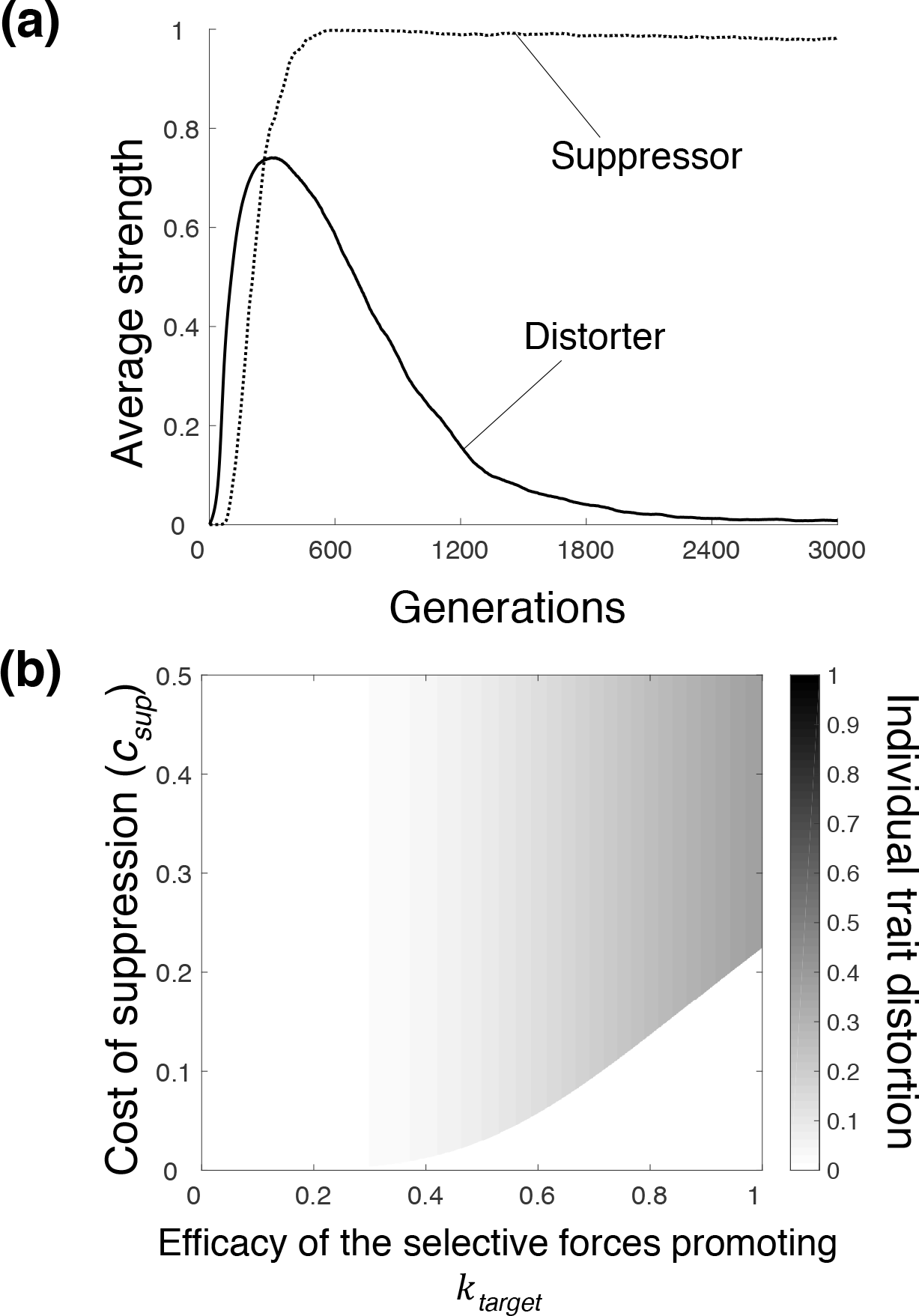
Evolution of trait distortion. In part (a), a distorter and suppressor are introduced in our agent-based simulation model (Appendix 7), and the following parameter regime is chosen: *c*_*sup*_=0.1,*t*=*k* and *c*_drive_ =max(*k_a_,k*_*b*_)/2. The population average distorter and suppressor strengths over 100 simulation runs are plotted for successive generations. Initially, both distorter and suppressor strength increases. Eventually, a threshold is passed, after which, distorters are purged from the population, meaning the trait is undistorted at equilibrium. In part (b), each location on the graph corresponds to a different parameter regime. Along the *x* axis, we vary the rate of increase in the marginal individual cost of trait distortion 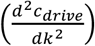 relative to the rate of decrease in the marginal transmission benefit 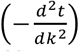, which determines the target level of trait distortion (*k*_target_) (Supplementary Information 1). Along the *y* axis, we vary the cost of suppression (*c*_*sup*_). We see that, when there is a low target level of distortion (left hand side of the heat map), suppressors fail to invade, leading to a low level of trait distortion at equilibrium. When there is a high target level of distortion and a relatively low cost of suppression (bottom right hand side), suppressors invade, leading to no trait distortion at equilibrium. When there is a high target level of distortion and a relatively high cost of suppression (top right hand side), suppressors do not invade, leading to appreciable trait distortion at equilibrium.

As evolution on the distorter increases the level of distortion, it makes it more likely that the distorter reaches the critical level of distortion where suppression will be favoured. When this is the case (*c_sup_<c*_*drive*_(*k*_*target*_)), the distorter spreads to high frequency, which then causes the suppressor to increase in frequency, reversing the direction of selection on the distorter, towards non-distortion (*y*_*0*_), resulting in zero trait distortion at equilibrium (*k*\*=0) (Fig. 2a; Appendix 6). Suppression only fails to spread if the individual fitness cost associated with suppression is greater than the individual fitness cost associated with the target trait distortion (*c_sup_>c*_*drive*_(*k*_*target*_); Fig. 2a). Given that the individual fitness cost of pre-translational suppression at a single locus is likely to be low, then any non-negligible distorter is likely to be suppressed.

Overall, our results suggest that selection on distorters will tend to drive the eventual suppression of those distorters. In Appendix 7 we developed an agent-based simulation, which allowed us to continuously vary the level of both distortion and suppression, and obtained results in close agreement (Fig. 2a; Supplementary Information 3, Fig. S2).

### Specific Models

We then tested the robustness of our above conclusions by developing models for three different biological scenarios: a sex ratio distorter on an X chromosome (X driver); an imprinted gene that is only expressed when maternally inherited; and a gene for the production of a public good by bacteria, which is encoded on a mobile genetic element. We examined these cases because they are different types of distortion, involving different selection pressures, in very different organisms. We obtained qualitatively similar results in all three cases.

In all of our specific models, we assume that the suppressor: is dominant; is only expressed in the presence of the distorter (facultative); completely suppresses the distorter; and may incur a fitness (viability) cost to the individual when it is expressed, independent of distorter strength, denoted by *c*_*sup*_ (0≤*c*_*sup*_≤1)^43,44^. These assumptions fit well to a molecularly characterised suppressor (“*nmy*”) of a sex ratio distorter (“*Dox”*)^39,40^; and more generally to suppressors that act pre-translationally^45,46^. We also relax a simplifying assumption of our illustrative model, by allowing the transmission benefit and individual fitness cost of trait distortion to vary with the population frequency of the distorter.

### Sex Ratio Distortion

We examined sex ratio evolution in a diploid species, in a large outbreeding (panmictic) population, with non-overlapping generations, and where males and females are equally costly to produce. Fisher^1^ and many others have shown that, in this scenario, individuals would be selected to invest equally in male and female offspring^9,24^. We assumed genetic sex determination, with males as XY and females as XX, and that females mate with λ mates per generation. The distorter (*y*_*1*_) that we considered is an X driving chromosome, which acts in males, killing Y-bearing sperm, and causing the male’s mating partners to produce a higher proportion of female (XX) offspring. The proportion is given by (1+*k*)/2, where *k* denotes the proportion of Y-bearing sperm that are killed (0<*k*≤1). We assumed that the sex ratio distorter can be suppressed by a costly autosomal suppressor (*sup*). This biology corresponds to sex ratio distortion in flies^25^.

Our sex ratio model showed very similar results to our illustrative model. In Supplementary Information 4 we show with population genetic analyses that when distortion is weak (low *k*), suppressors are not favoured. In contrast, if distortion is strong (high *k*), then suppression is favoured, and so there is no net influence on the individual trait value. Consequently, the extent to which the sex ratio deviates from a 50:50 investment will be small or zero^47^ (Fig. 3ai). In addition to being more likely to be suppressed, the stronger distortion is, the quicker suppressors spread (Fig. S5). Finally, when we allowed the X chromosome driver to evolve, it evolved to high levels of sex ratio distortion (high *k*_*target*_), increasing the likelihood of suppression.

**Figure 3.**
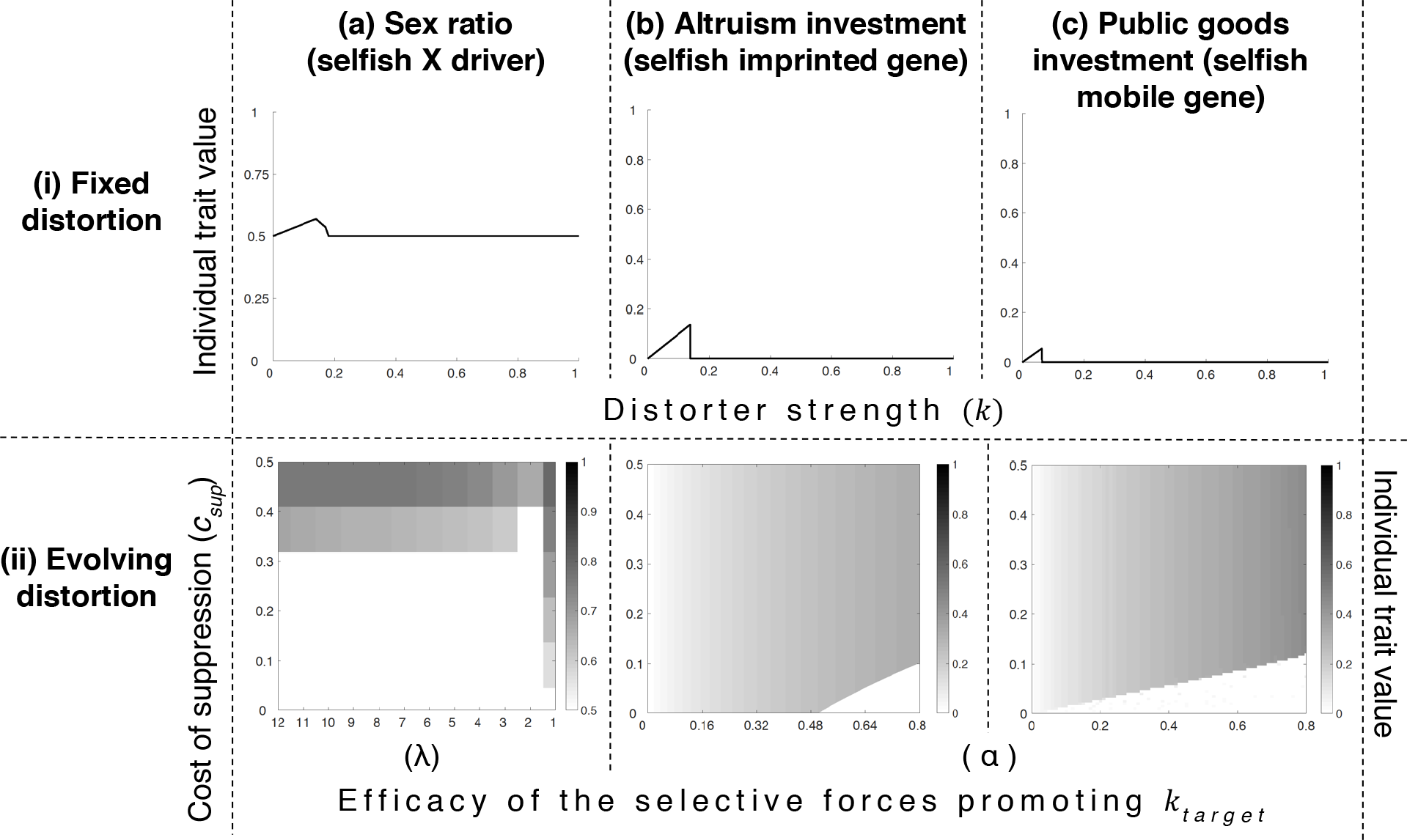
Specific Biological Scenarios. We consider three biological scenarios: (a) sex ratio distortion by an X driver; (b) cooperative investment by an imprinted gene, in which the cost of cooperation is assumed to be *c*=*k* and the benefit is *b*=*k*^a^; (c) cooperative public goods investment by a mobile gene, in which the cost of cooperation is assumed to be *c*=*k* and the benefit is *b*=8*k*^a^. Part (i) plots equilibrium trait values after short term coevolution between distorters of different strengths (*k*) and their suppressors. In part (i), the cost of suppression is taken to be *c*_sup_ =0.05; double female mating (λ=2) is assumed in (ai); and slightly decelerating returns on cooperation (α=0.9) is assumed in (bi) and (ci). Part (i) shows that trait distortion will be relatively low or absent. Part (ii) shows the level of distortion that will be evolved to, when distorters can evolve. Along the x axis, we vary a model parameter that affects the target level of trait distortion (*k*_target_). In (aii), we vary the female mating rate (λ), and in (bii) and (cii), we vary α, which mediates the rate of decrease in marginal cooperative benefits 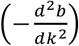 relative to the rate of increase in marginal cooperative costs 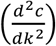. Part (ii) demonstrates that trait distortion will evolve to be zero or relatively low, except when there is a high cost of suppression (*c*_sup_) alongside a high target level of distortion (low λ / high α), corresponding to the top right regions of the heat maps.

We only obtained appreciable and detectable levels of sex ratio distortion (>60% females) if the cost of suppression exceeded a 15-35% viability reduction (Fig. S8). This is a much greater cost than what we would expect from natural gene suppression pathways^48^. A suppressor will only fail to spread when the individual cost of sex ratio distortion is less than the cost of suppressing the distorter (Equation S3). Given that the cost of suppression is likely to be low, we would only expect distorters that have relatively little impact at the individual level to evade suppression. We tested the robustness of our population genetic analysis with an agent-based simulation, and a game theory model, and found close agreement (Fig. S6).

Our predictions are consistent with the available data on X drivers in *Drosophila*. As predicted by our model: (1) Across natural populations of *D. simulans*, there is a positive correlation between the extent of sex ratio distortion and the extent of suppression^49^. (2) In both *D. mediopunctata* and *D. simulans* the presence of an X linked driver led to the experimental evolution of suppression^50,51^. In addition, consistent with our model: (3) In natural populations of *D. simulans* the prevalence of an X driver has been shown to sometimes decrease under complete suppression^52^. (4) Crossing different species of *Drosophila* has been shown to lead to appreciable sex ratio deviation, by unlinking distorters from their suppressors, and hence revealing previously hidden distorters^53^. Work on other sex ratio distorters has also shown that suppressors can spread extremely quickly from rarity, reaching fixation in as little as ~5 generations^54^.

### Genomic Imprinting and Altruism

Genomic imprinting occurs at a minority of genes in mammals and flowering plants. An imprinted allele has different epigenetic marks, and corresponding expression levels, when maternally and paternally inherited^55^. We examined the evolution of an altruistic helping behaviour in a population capable of genomic imprinting. A behaviour is altruistic if it incurs a cost (*c*) to perform, by the actor, and provides a benefit (*b*) to another individual, the recipient. Altruism is favoured if the genetic relatedness (*R*) between the actor and recipient is sufficiently high, such that *Rb*>*c*^3^.

An individual may be more closely related to their social partners via their maternal or paternal genes^14,15,18^. For example, if a female mates two males, then on average her offspring would be related by *R_m_*=1/2 at maternal genes and *R*_*p*_=1/4 at paternal genes. If genes can ‘gain information’ about where they came from, by imprinting, then they could be selected to adjust traits accordingly. Assume that relatedness to social partners is *R*_*p*_ and *R*_*m*_ at paternal and maternal genes respectively. In this case, altruistic helping would be favoured at: maternally imprinted genes when *R_m_b*>*c*; paternally imprinted genes when *R_p_b*>*c*; and unimprinted genes when ((*R*_*p*_+*R*_*m*_)/2*)b*>*c*^18,56,57^. Consequently, if *R_m_b*>*c*>((*R*_*p*_+*R*_*m*_)/2)*b*, then altruistic helping is favoured at maternally imprinted genes, when it is disfavoured at unimprinted genes (selfish trait distortion).

We modelled the evolution of altruism in a large population of diploid, sexually reproducing individuals. The distorter (*y*_*1*_) increases altruistic investment by some amount (*k*), at a fitness cost to the individual (0≤*c*_*GI*_(*k*)≤1) and benefit to the social partner (*bGI*(*k*)>*c*_*GI*_(*k*)) that are both monotonically increasing functions of investment 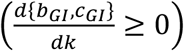. The distorter (*y*_1_) is only expressed when maternally inherited, whereas its potential suppressor (*sup*) is unimprinted^58^. Every generation, individuals associate in pairs with kin that they are maximally related to via their maternally inherited genes (*R*_*m*_=1) but minimally related to via their paternally inherited genes (*R*_*p*_=0). Individuals then have the opportunity to be altruistic to their partner, before mating at random in proportion to their fitness (fecundity), reproducing, then dying (non-overlapping generations).

In Supplementary Information 5, we showed with a population genetic analysis that our imprinting model produces very similar results to our illustrative model. When distortion is weak (low *k*), such that the cost of trait distortion is less than the cost of suppression, suppressors are not favoured (Equation S8). Given that the distorter suppression cost is likely to be low, these distorters that evade suppression will have relatively little impact at the individual level. When distortion is strong (high *k*), then suppression is favoured, and so there is no influence on the individual trait value^58^. Consequently, the extent to which altruistic investment deviates from the individual optimum of zero investment will be small or zero (Fig. 3bi). Finally, when we allowed the imprinted gene to evolve, we found that cooperative distortion increased until it reached the value (*k*_*target*_) at which the marginal benefit of cooperation is cancelled out by the marginal cost of cooperation 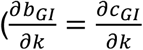; Equation S10), increasing the likelihood of suppression (Fig. 3bii).

Although there have been no direct tests, our predictions are consistent with data on imprinted genes. There is no evidence that traits influenced by imprinted genes deviate significantly from individual level optima under normal development^14^. Significant deviation is only observed when imprinted genes are deleted, implying that imprinted trait distorters are either suppressed, or counterbalanced by oppositely imprinted genes pulling the trait in the opposite direction^32,58^. Furthermore, although many different parties (coreplicons) have vested interests in genomic imprinting, our analysis suggests why the unimprinted majority could win control^59^. This could help explain both why imprinting appears to be relatively rare within the genome^18,55,60^, and why imprints are removed and re-added every generation in mice, handing control of genomic patterns of imprinting to unimprinted genes^18,59,61^.

### Horizontal Gene Transfer and Public Goods

Bacteria produce and excrete many extracellular factors that provide a benefit to the local population of cells and so can be thought of as public goods^62^. We modelled the evolution of investment in a public good in a large, clonally reproducing population. We assume a public good that costs *c* to produce, and provides a benefit *b* to the group. We assume a well-mixed population, meaning genetic relatedness at vertically inherited genes is zero (*R*_*vertical*_=0), and so public good production is disfavoured at the individual level (*R_vertical_b*=0<*c*)^3,63^.

We consider a distorter (*y*_*1*_) of public goods production that is on a mobile gene, such as a plasmid. Mobile genes can spread within groups, increasing genetic relatedness at the mobile locus (*R*_*horizontal*_>0), potentially favouring public goods production^64,65^. We assume that the distorter increases public goods investment by some amount (*k*), at a fitness cost to the individual (0≤*c*_*HGT*_(*k*)≤1) and benefit shared within the group (*b*_*HGT*_(*k*)>*c*_*HGT*_(*k*)), that are both monotonically increasing functions of investment 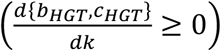. We also assume a potential suppressor (*sup*) that is immobile^46,66,67^. Each generation, individuals randomly aggregate into groups, and one allele at the mobile locus (*y*_0_,*y*_1_,*y*_2_) spreads horizontally within each group, each with equal likelihood, increasing relatedness at the mobile locus. Public goods may then be produced and shared within groups, before individuals reproduce and die (non-overlapping generations).

In Supplementary Information 6, we show with a population genetic analysis that our plasmid model produces similar results to our illustrative model. When distortion is weak (low *k*), suppressors are not favoured, but the distorter has relatively little impact at the individual level (Fig. 3ci). When distortion is strong (high *k*), then suppression is favoured, and so there is no influence on the individual trait value^67^. Consequently, the extent to which public goods investment deviates from the individual optimum of zero investment will be relatively small or zero (Fig. 3ci). Finally, when we allowed the mobile distorter to evolve, we found that higher levels of public goods investment (high *k*_*target*_) are favoured, leading to selection for suppression (Fig. 3cii). We lack empirical data that would allow us to test our model of mobile public goods genes.

## Discussion

We have found that the individual level consequences of selfish genetic elements (‘*distorters*’) will be either small or non-existent. If distorters lead to only small distortions of traits, then they will not be suppressed, but they will only have small effects on traits (Figs. 1a & 3i). Specifically, distorters will only remain unsuppressed if trait distortion compromises individual fitness less than suppression of the trait distorter does (*c*_*drive*_(*k*)<*c*_*sup*_). Given that the individual fitness cost of pre-translational suppression at a single locus is likely to be low, we can say that trait distortion conferred by unsuppressed distorters is likely to be relatively negligible.

However, if distorters lead to large distortions of traits then this selects for their suppression, and so they will have no net effect on traits at the individual level (Figs. 1a & 3i). Stronger distorters will also be suppressed more quickly (Fig. 1b). Furthermore, selection on distorters favours higher levels of distortion, which will render them more likely to be ultimately suppressed. Consequently, the evolution of distorters will often drive their own demise (Figs. 2, 3ii & 4). These results suggest that even if there is substantial potential for genetic conflict, distorters will have relatively little influence at the individual level, in support of Leigh’s^27^ parliament of genes hypothesis.

**Figure 4:**
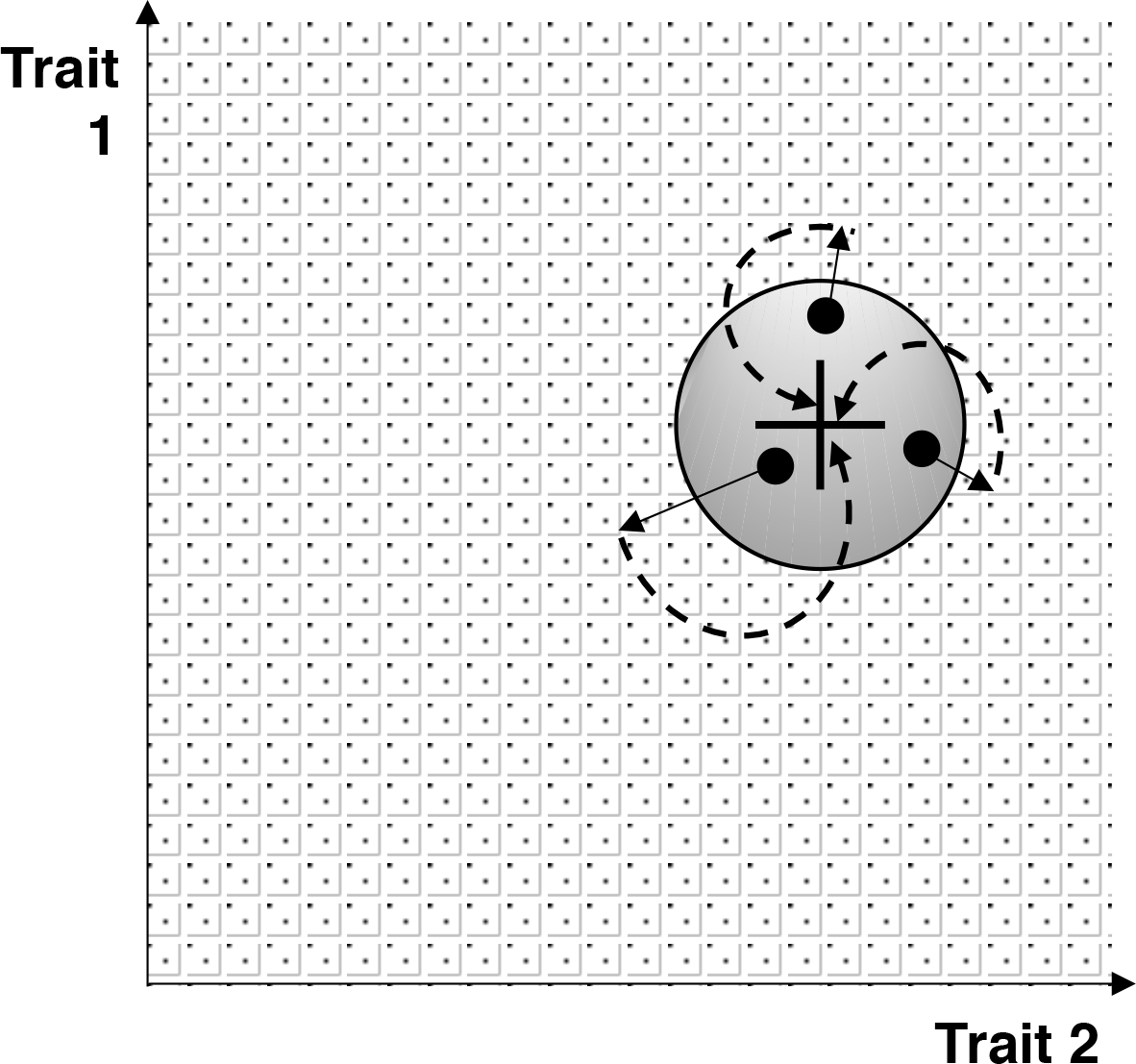
Selfish genetic elements evolve to be suppressed by the parliament of genes. The cross represents the position in phenotype space, here defined with respect to two traits, *1* and *2*, that maximises the fitness of an individual. The circle surrounding the cross represents the phenotype space where suppression of selfish genetic elements, that have distorted traits 1 or 2, would not be selected for. The surrounding area represents the phenotype space in which the parliament of genes is selected to suppress selfish genetic elements. The three dots represent three possible individuals, which, owing to weakly selfish genetic elements, are not maximising individual fitness (the dots do not lie exactly on the cross), although they are approximately (e.g. within the bounds of experimental error in measurement). Because these deviations from individual fitness maximisation are only slight, costly suppression of the weakly selfish genetic elements does not evolve. However, the selfish genetic elements will evolve to become more distorting (solid arrows), bringing individuals into the non-tolerated area of phenotype space, where they will be suppressed and individual fitness maximisation (the black cross) is regained (dashed arrows).

We have assumed that there will be a much greater number of genes where suppression is favoured, and so it is relatively easy for a suppressor to be reached by mutation^31^. We have therefore examined, given the potential for suppression, what direction would we expect natural selection to take on average. We are not claiming that appreciable trait distortion will never evolve, and there are clearly cases where it has^14^. A number of biological details will matter for different systems, including: (i) whether a cost of suppression is incurred when the distorter is not present; (ii) whether distorters can outstrip suppressors in coevolutionary arms races, leading to distorters going unsuppressed for extended time periods^31,54,68^; (iii) the rate at which trait-distorting selfish genetic elements arise by mutation, relative to the rate at which their suppressors arise by mutation. Our results are supported by cases where appreciable distortion is only revealed in hybrid crosses, implying that they are generally suppressed^53,69^.

We emphasise that when the assumption of individual fitness maximisation is made in behavioural and evolutionary ecology, it is not being assumed that natural selection produces perfect fitness maximisers^5^. Many factors could constrain adaptation, such as genetic architecture, mutation and phylogenetic constraints^70,71^. Instead, the assumption of fitness maximisation is used as a basis to investigate the selective forces that have favoured particular traits (adaptations). The aim is not to test if organisms maximise fitness, or behave ‘optimally’, but rather to try to understand the selective forces favouring particular traits or behaviours^2^.

To conclude, debate over the validity of assuming individual level fitness maximisation has revolved around whether selfish genetic elements are common or rare^4,19,20,23,72^. We have shown that that even if selfish genetic elements are common, they will tend to be either weak and negligible, or suppressed. This suggests that even if there is the potential for appreciable genetic conflict, individual level fitness maximisation will still often be a reasonable assumption. This allows us to explain why certain traits, especially the sex ratio, have been able to provide such clear support for both individual level fitness maximisation and genetic conflict.

## Appendix

## Appendix 1: Distorter population frequency

We ask when a rare distorter (*y*_1_) can invade a population fixed for the non-distorter (*y*_0_). We take Equation (1), set *p*’=*p*=*p*\*, and solve to find two possible equilibria: *p\**=0 (non-distorter fixation) and *p\**=1 (distorter fixation). The distorter (*y*_1_) can invade from rarity when the *p*\*=0 equilibrium is unstable, which occurs when the differential of *p*’ with respect to *p*, at *p*\*=0, is greater than one. The distorter invasion criterion is therefore *c*_*drive*_(*k*)<*t*(*k*)(1*-c*_*drive*_(*k*)).

We now ask what frequency the distorter (y_1_) will reach after invasion. The distorter (*y*_1_) can spread to fixation if the *p*\*=1 equilibrium is stable, which requires that the differential of *p*’ with respect to *p*, at p*=1, is less than one. This requirement always holds true, demonstrating that there is no negative frequency dependence on the distorter, and that it will always spread to fixation after its initial invasion.

## Appendix 2: Suppressor invasion condition

We ask when the suppressor (*sup*) can spread from rarity in a population in which the distorter (*y*_1_) and non-suppressor (+) are fixed at equilibrium. We derive the Jacobian stability matrix for this equilibrium, which is a matrix of each genotype frequency (*x*_1_’, *x*_2_’, *x*_3_’, *x*_4_’) differentiated by each genotype frequency in the prior generation (*x*_1_, *x*_2_, *x*_3_, *x*_4_), at the equilibrium position given by *x*_*1*_*=0, *x*_*2*_*=0, *x*_*3*_*=1, *x*_*4*_*=0:

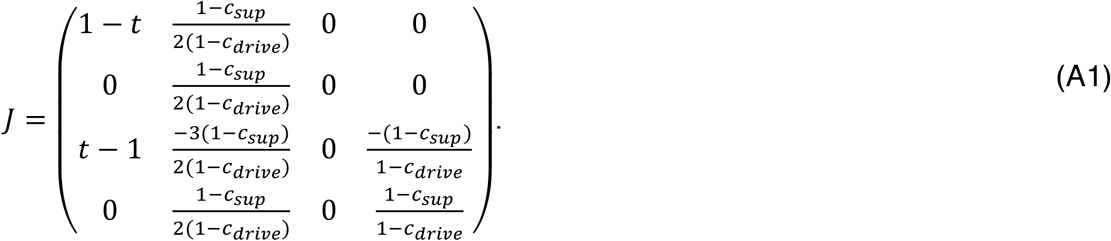

The suppressor can invade when the equilibrium is unstable, which occurs when the leading eigenvalue is greater than one. The leading eigenvalue is (1-*c*_*sup*_)/(1-*c*_*drive*_), meaning the suppressor invasion criterion is *c*_*drive*_>*c*_*sup*_.

## Appendix 3: Equilibrium distorter and suppressor frequencies

We ask what frequency the distorter (*y*_1_) and suppressor (*sup*) will reach after initial suppressor (*sup*) invasion. We assume that the suppressor is introduced from rarity when the distorter has reached the population frequency given by *f* (*x*_1_→*f*, *x*_3_→1-*f*, {*x*_2_,*x*_4_}→0). We numerically iterate Equations (2), over successive generations, until equilibrium has been reached. At equilibrium, for all parameter combinations (*f*,*t,c*_*sup*_,*c*_*drive*_), the suppressor reaches an internal equilibrium and the distorter is lost from the population (*x*_*1*_*+*x*_*2*_*=1, *x*_*3*_*=0, *x*_*4*_*=0). This equilibrium arises because distorter-presence gives the suppressor (*sup*) a selective advantage, leading to high suppressor frequency, which in turn reverses the selective advantage of the distorter (*y*_1_), leading to distorter loss and suppressor equilibration.

## Appendix 4: Non-equilibrium trait distortion

We consider a distorter that is suppressed and therefore purged at equilibrium (*c*_*drive*_>*c*_*sup*_), and ask to what extent it can contribute to individual trait distortion in the period after its initial invasion but before its eventual loss (non-equilibrium). We introduce the distorter (*y*_1_) and suppressor (*sup*) from rarity and numerically iterate our recursions until the distorter has been purged from the population (or a cap of 20,000,000 generations has been reached). We vary parameters between 0≤*t*≤1, *c*_*sup*_<*c*_*drive*_≤1, 0≤*c*_*sup*_≤1.

We find that a higher cost of trait distortion (*c*_*drive*_) relative to suppression (*c*_*sup*_) leads to shorter non-equilibrium maintenance of the distorter in the population. This is because the cost of trait distortion relative to suppression mediates selection on the suppressor (Appendix 2). We find that a higher transmission bias (*t*) leads to longer non-equilibrium maintenance of the distorter in the population, but this effect is diluted as the cost of trait distortion (*c*_*drive*_) is increased relative to suppression (*c*_*sup*_) (Supplementary Information 2, Fig. S1). Stronger distorters (with higher *k*, leading to higher *c*_*drive*_ and *t*) are therefore generally suppressed and purged more rapidly than weaker distorters (Fig. 1b). Exceptions are distorters that reduce individual fitness relatively negligibly after the point (*k*) at which suppression is favoured, such that 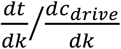 is very high for values of *k* satisfying *c_sup_<c*_*drive*_(*k*).

## Appendix 5: Invasion of a mutant distorter

We ask when a mutant distorter (*y*_2_) will invade against a resident distorter (*y*_1_) that is unsuppressed and at fixation 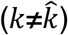. We write recursions detailing the generational frequency changes in the six possible gametes, *y_0_*/+, *y_0_*/*sup*, *y_1_*/+, *y_1_*/*sup*, *y_2_*/+, *y_2_*/*sup*, with current generation frequencies denoted respectively by *x*_*1*_, *x*_*2*_, *x*_*3*_, *x*_*4*_, *x*_*5*_, *x*_*6*_, and next generation frequencies denoted with an appended dash (’):

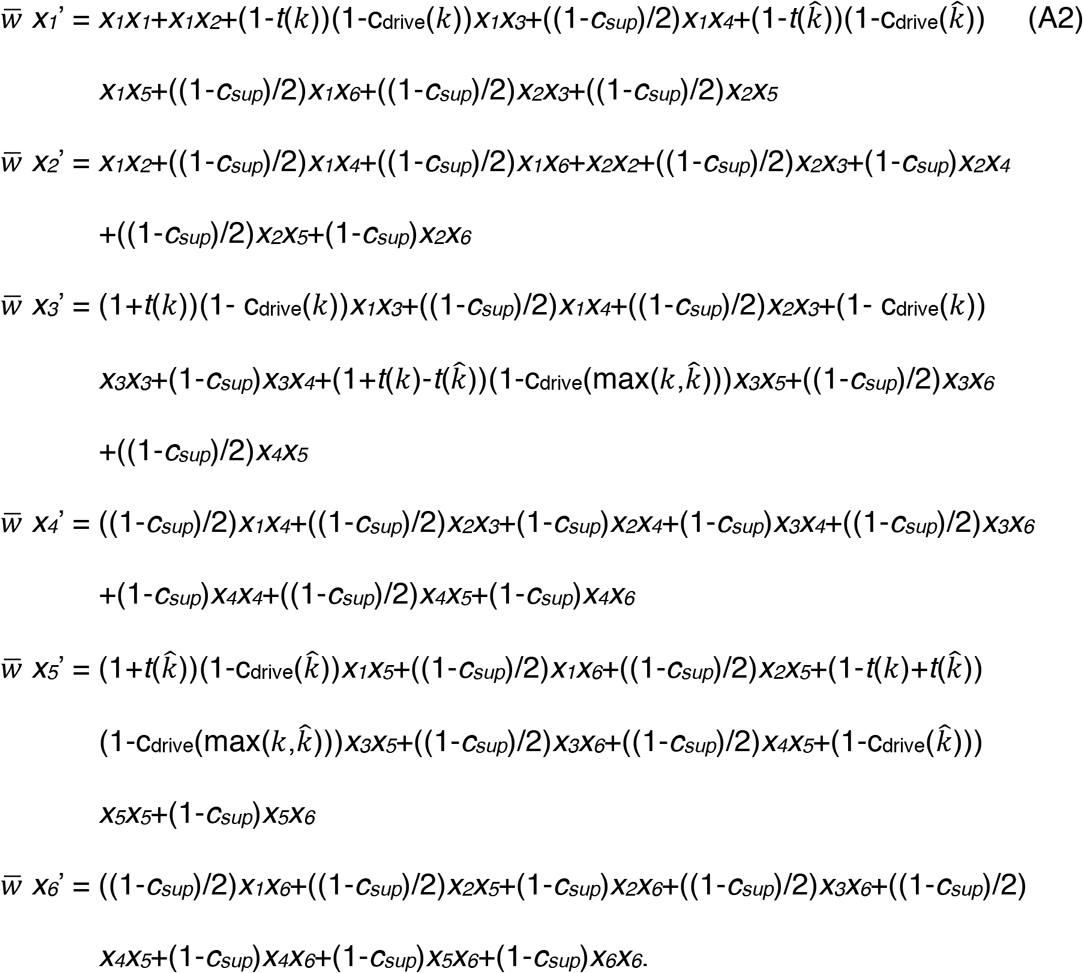

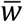 is the average fitness of individuals in the current generation, and equals the sum of the right-hand side of the system of equations. The mutant distorter can invade when the equilibrium given by *x*_*1*_*=0, *x*_*2*_*=0, *x*_*3*_*=1, *x*_*4*_*=0, *x*_*5*_*=0, *x*_*6*_*=0 is unstable, which occurs when the leading eigenvalue of the Jacobian stability matrix for this equilibrium is greater than one. Testing for stability in this way, we find that, if the mutant distorter is weaker than the resident, it can never invade. If the mutant distorter is stronger than the resident, it invades from rarity when 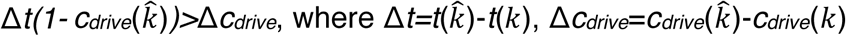.

The implication is that, if trait distortion is initially low, and mutant distorters are successively introduced, each deviating only very slightly from the resident distorter from which they are derived, such that 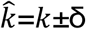, where δ is very small (“δ-weak selection”^42^), then distorters will approach a ‘target’ strength at which 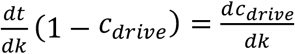. In the absence of suppression, this target (*k*_*target*_) is the equilibrium level of distortion (*k*\*=*k*_*target*_). However, if mutant distorters (*y*_2_) are allowed to deviate appreciably from residents (*y*_1_) (strong selection), then distorters may invade even if they overshoot the target 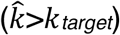. In the absence of suppression, *k*_*target*_ is then not the equilibrium level of distortion, but rather, the minimum equilibrium level of distortion (*k*\*>*k*_*target*_) (Supplementary Information 3, Fig. S2b).

We could alternatively have assumed that an individual’s trait is distorted according to the average strength of its alleles (additive gene interactions), rather than according to the stronger (higher-*k*) allele (dominance). Such an assumption leads to a single invasion criterion for a mutant distorter, regardless of whether the mutant distorter is stronger or weaker than the resident distorter, given by: 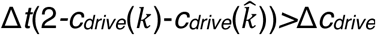. In the absence of suppression, this leads to an equilibrium level of distortion (*k*\*), that holds even under strong selection, that satisfies 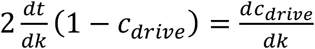.

## Appendix 6: Equilibrium distorter and suppressor frequencies (long term evolution)

We ask what equilibrium state will arise after the invasion of a mutant distorter. We assume that the mutant distorter (*y*_2_) is introduced from rarity when the resident distorter (*y*_1_) has reached the population frequency given by *q*. We numerically iterate Equations (A2), over successive generations, until equilibrium has been reached. At equilibrium, for all parameter combinations 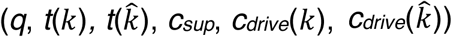, the resident distorter (*y*_*1*_) is lost from the population (*x*_*3*_,*x*_*4*_=0), with either the mutant distorter (*y*_2_) and non-suppressor (+) at fixation (*x*_*5*_*=1), or the non-distorter at fixation alongside the suppressor at an internal equilibrium (*x*_*1*_*+*x*_*2*_*=1). The latter scenario arises if the mutant distorter triggers suppressor invasion 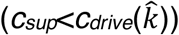. This equilibrium arises because mutant distorter-presence gives the suppressor (*sup*) a selective advantage, leading to high suppressor frequency, which in turn reverses the selective advantage of distortion, leading to distorter (*y*_1_,*y*_2_) loss and suppressor equilibration.

## Appendix 7: Agent-based simulation

We construct an agent-based simulation to ask what level of trait distortion evolves when continuous variation is permitted at distorter and suppressor loci. We model a population of N=2000 individuals and track evolution at two autosomal loci: a distorter locus (L1) and a suppressor locus (L2). Each individual has two alleles at the distorter locus, with strengths denoted by *k_a_* and *k_b_*, and two alleles at the suppressor locus, with strengths denoted by *m_a_* and *m_b_* (diploid). Strengths can take any continuous value between zero and one. We assume that, for both loci, the strongest (highest value) allele within an individual is dominant. The absolute fitness of an individual with at least one active meiotic driver (max(*k_a_*,*k_b_*)>0) is: 1-*c_drive_*(max(*k_a_*,*k_b_*))(1-max(*m_a_*,*m_b_*))-*c_sup_*max(*m_a_*,*m*_*b*_), and the absolute fitness of an individual lacking an active distorter (max(*k*_*a*_,*k*_*b*_)=0) is 1. The function *c*_*drive*_(max(*k*_*a*_,*k*_*b*_)) is given an explicit form in simulations (Supplementary Information 3, Fig. S2).

Each generation, there are N breeding pairs. To fill each position in each breeding pair, individuals are drawn from the population, with replacement, with probabilities given by their fitness (hermaphrodites). Breeding pairs then reproduce to produce one offspring, before dying (non-overlapping generations). Alleles at the suppressor locus (L2) are inherited in Mendelian fashion. Alleles at the distorter locus may drive, meaning the parental allele of strength *k_a_* is inherited, rather than the allele of strength *k*_*b*_, with the probability (1+(*t*(*k*_*a*_)-*t*(*k*_*b*_))(1-max(*m*_*a*_,*m*_*b*_)))/2. The transmission bias function, *t*, is given an explicit form in simulations (Supplementary Information 3, Fig. S2). Each generation, distorter and suppressor alleles have a 0.01 chance of mutating to a new value, which is drawn from a normal distribution centred around the pre-mutation value, with variance 0.2, and truncated between 0 and 1. We track the population average distorter strength, denoted by E[*k*], and suppressor strength, denoted by E[*m*], over 20,000 generations. We see that, allowing for continuous variation at the distorter and suppressor loci, if the cost of suppression (*c*_*sup*_) is not excessively high, trait distortion at equilibrium is either low or nothing (Fig. 2a; Fig. S2b).

## Acknowledgements

We thank Geoff Wild, Alan Grafen, Egbert Giles Leigh, Jr., Guy Cooper, Sam Levin, Asher Leeks and Matishalin Patel for helpful comments and discussion.

## Author Contributions

TWS and SAW designed the study and wrote the paper. TWS carried out mathematical analysis.

## Competing Interests

The authors declare no competing interests.

## Data Accessibility

The data that support the findings of this study are available from the corresponding author upon reasonable request.

