## Supplementary Information 1-7 for "Adaptation and the Parliament of Genes"


(1) Functional forms assumed in Fig. 2b (Main Text) p1

(2) Figure S1 p2
Non-equilibrium trait distortion

(3) Figure S2 p3
Comparison of population genetic and agent based simulation results of the illustrative model

(4) Sex Ratio Distortion p4

(5) Genomic Imprinting and Altruism p28

(6) Horizontal Gene Transfer and Public Goods p39

(7) Additional Discussion p47
- Suppressors
- Dynamics and Mutation
- Individual Fitness Maximisation and the Parliament of Genes

References p49

### 1) Functional forms assumed in Fig. 2b (Main Text)

To generate Figure 2b (Main Text), we assumed the functional forms $c_{drive}$=0.8$k^{1.3}$ and $t$=T1$k$, where 0≤T1≤1. The parameter T1 mediates the rate at which the marginal transmission advantage dissipates, relative to the marginal individual cost of trait distortion, as the trait becomes increasingly distorted ($k$). This parameter (T1) is plotted along the *x* axis of Figure 2b (Main Text). We analytically derived the target trait distortion ($k_{target}$), for different values of $c_{sup}$ and T1, by substituting our specific functional forms into the condition that specifies $k_{\text{target}}$: $\frac{dt}{dk}(1 - c_{drive}) = \frac{dc_{drive}}{dk}$. This gave: $\frac{d(T1\ k)}{dk}(1 - 0.8k^{1.3}) = \frac{d(0.8k^{1.3})}{dk}$, which simplifies to T1(1-0.8$k^{1.3}$=1.04$k^{0.3}$), which was solved for $k$ to give $k = k_{target}$. We derived the equilibrium trait distortion ($k^{*}$) by substituting each value of $k_{target}$ into the condition for suppressor spread: $c_{sup}$<$c_{drive}$($k_{target}$). Satisfaction of this condition implies that $k^{*}$=0; else, $k^{*}$=$k_{target}$.

### 2) Figure S1

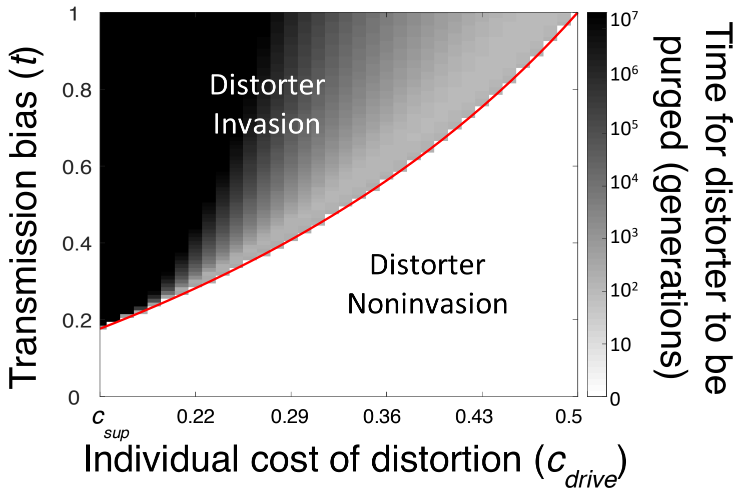


**Figure S1. Non-equilibrium trait distortion.** A trait distorter ($y_1$) is introduced from rarity alongside its suppressor (*sup*) (initial genotype frequencies: $x_1$=0.97, {$x_2$,$x_3$,$x_4$}=0.01). The trait distorter ($y_1$) is associated with some individual cost when expressed ($c_{drive}$) which is varied along the x axis (the cost of suppression is fixed at $c_{sup}$=0.15). The trait distorter ($y_1$) is also associated with a transmission bias at meiosis ($t$) which is varied along the y axis. We consider distorters that induce suppressor spread ($c_{sup}<c_{drive}$) and ask whether such distorters can cause appreciable trait distortion before they are ultimately suppressed and purged from the population. The red line plots the formula $t=c_{drive}/(1-c_{drive})$; above this line, distorters can spread from rarity. We plot the number of generations (on a $\log_{10}$ scale) until equilibrium is reached (trials that did not equilibrate by 20,000,000 generations were capped). We see that less costly distorters ($c_{drive}$ only slightly greater than $c_{sup}$) can invade even with a relatively low transmission bias ($t$), and are purged at a very slow rate, causing extended non-equilibrium trait distortion. More costly distorters ($c_{drive}$ large compared to $c_{sup}$) require a high transmission bias (*t)* to invade, and if they can invade, they are purged relatively quickly, causing shorter non-equilibrium trait distortion. Therefore, non-equilibrium trait distortion is either not-so-costly and extended, or costly and ephemeral, and so has limited impact on individual fitness maximisation in either case.

### 3) Figure S2

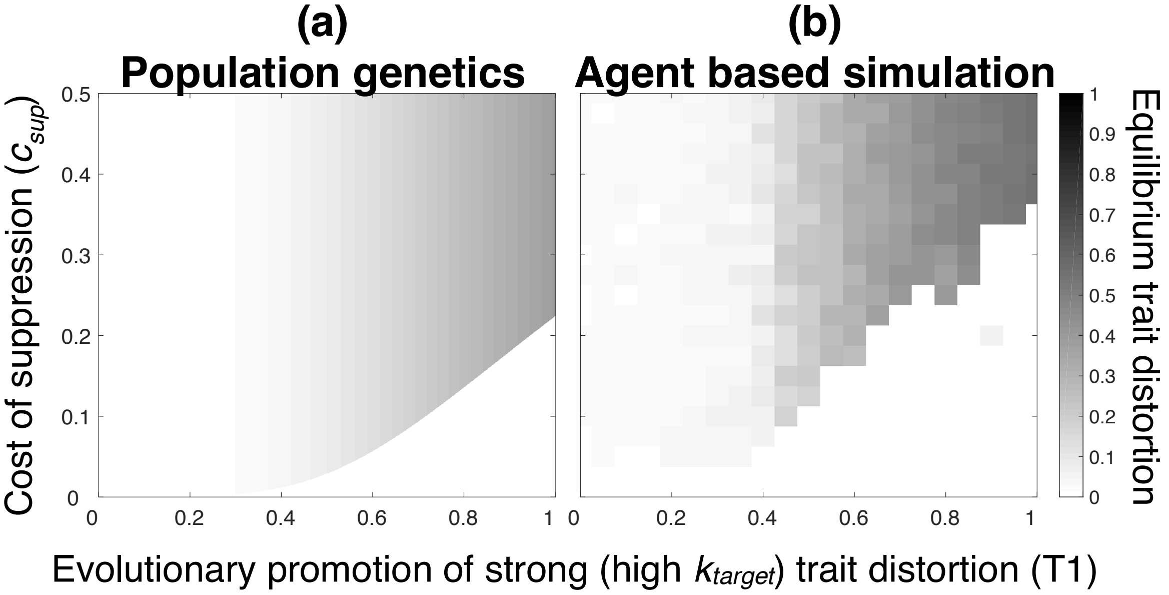


**Figure S2. Comparison of population genetic and agent based simulation results of the illustrative model.** Distorter strength evolves in the presence of a suppressor of distortion (*sup*) and the resulting equilibrium trait distortion ($0 \leq k^* \leq 1$) is plotted, using the following functional forms for distorter cost ($c_{drive}$) and transmission advantage ($t$): $c_{drive}(k) = 0.8k^{1.3}$ and $t(k) = T1k$, where $0 \leq T1 \leq 1$, and where T1 and $c_{sup}$ are varied. T1 mediates the rate at which the marginal transmission advantage $\left(\frac{dt}{dk}\right)$ dissipates, relative to the marginal individual cost $\left(\frac{dc_{drive}}{dk}\right)$ of trait distortion, as the trait becomes increasingly distorted (*k*). Part (a) plots the equilibrium trait distortion under weak selection, as analytically derived from the population genetic model in the main text. Part (b) plots the equilibrium trait distortion as obtained by the agent based simulation detailed in Appendix 7, which does not assume weak selection, and which allows continuous variation at the trait and suppressor loci. There is good correspondence between the models; however, the region in which distorters are suppressed ($k^*=0$; bottom right) is larger in the agent based simulation, and outside of this region, the equilibrium trait distortion ($k^*$) is slightly greater in the agent based simulation. Discrepancy arises from the different assumptions about the strength of selection.

### 4) Sex Ratio Distortion

We consider a selfish genetic element residing on an X chromosome, that may gain a propagation advantage by distorting the offspring sex ratio towards a greater production of females. The genes that do not gain a propagation advantage from female sex ratio bias reside on both the autosomes and the Y chromosome[1]. We focus on suppressors in the autosomes, for simplicity, and because this is the larger group of genes (*coreplicon*[2]), constituting the majority within the parliament of genes. Consequently, we focus our analyses on when an X driver and an autosomal suppressor can spread.

Our overall aim is to assess, given the potential for suppression, the extent that an X chromosome driver can distort the sex ratio away from the individual optimum. The individual optimum is taken to be the evolutionarily stable strategy (ESS) adopted by individuals in the absence of selfish trait distortion, which is an equal investment in offspring of both sexes.

We build our model in a step-wise manner, as described in the “Results” section of the main text. Aspects of questions 1-3 have been analysed before with respect to sex ratio, but we go over them here for the specific case of our model, and to elucidate the underlying selective forces. There is available data on the fitness consequences of sex ratio distortion and suppression, and so, in this case, we aim for a biologically realistic model that can be parameterised.

#### (1) Spread of a Distorter

We considered the spread, in the absence of suppression, of a selfish sex ratio distorter that skews offspring sex ratio towards females. In the literature, selfish X drivers are often denoted by *SR* (for *sex-ratio* distortion), with non-distorting rival alleles denoted by *ST* (for *standard*)[3]. However, we denote the distorting and non-distorting alleles respectively by $y_1$ and $y_0$ for consistency across our models. We assume that normal ($y_0/Y$) males produce X and Y sperm equally. The distorter ($y_1$) causes $y_1/Y$ males to kill Y-bearing sperm, leading to a female-biased sex ratio[4-9]. In males with an unsuppressed distorter, its proportion of X-bearing sperm, and correspondingly, the proportion of its offspring that are female, is given by $0.5(1+k)$, where $k$ denotes the proportion of Y-bearing sperm that are killed ($0<k\leq 1$).

We assume that males with an unsuppressed distorter ($y_1$) suffer a fertility cost as a result of sperm death, and have a reduced ejaculate size of $1-k/2$, relative to 1 in all other males[10-15]. We assume that, each generation, each female copulates with λ random males, and that each sperm cell is equally competitive in the female's internal store. The likelihood of a male's sperm fertilising an egg (*fertility, F*) is given by his ejaculate size relative to the total amount of ejaculate that female has received. Letting $l$ be the proportion of males in the present generation with an unsuppressed distorter, the fertility of those males with an unsuppressed distorter ($F_{drive}$), and the fertility of those without ($F_{normal}$), is given by:

$$F_{drive} = \sum_{i=1}^{\lambda} \frac{\left(1-\frac{k}{2}\right)}{\left(1-\frac{k}{2}\right)i+(\lambda-i)} l^{i-1}(1-l)^{\lambda-i}, \quad \text{(S1a)}$$

$$F_{normal} = \sum_{i=1}^{\lambda} \frac{1}{\left(1-\frac{k}{2}\right)(i-1)+(\lambda-i+1)} l^{i-1}(1-l)^{\lambda-i}. \quad \text{(S1b)}$$

There is no sperm competition, and therefore no fertility cost of sex ratio distortion, when females are singly mated ($F_{normal} = F_{drive}$ when λ=1). There is increased sperm competition at higher female mating rates, meaning the relative fertility cost of sex ratio distortion ($F_{normal}/F_{drive}$) increases and plateaus for high λ at $F_{normal}/F_{drive} = 1/(1-k/2)$.

The distorter has no fitness consequences for females, and so the condition for the spread of the distorter ($y_1$) allele is that $y_1/Y$ males sire more female offspring than $y_0/Y$ males. In the absence of suppression, $y_1/Y$ males have $F_{drive}(1+k)/2$ female offspring, and $y_0/Y$ males have $F_{normal}/2$ female offspring, meaning the distorter ($y_1$) is selected when:

$$F_{normal} / F_{drive} < (1+k). \qquad \text{(S2)}$$

The left-hand side of Equation S2 gives the between-individual relative fertility cost of trait distortion and the right-hand side gives the within-individual relative transmission advantage of trait distortion. When we substitute our explicit fertility functions (Equation S1) into Equation S2, we find that Equation S2 is always satisfied. Consequently, analogous to previous arguments, the distorting $y_1$ chromosome will always spread to fixation, irrespective of female mating frequency (λ)[16,17]. This distorts the offspring sex ratio, defined as the proportion of females, to $(1+k)/2$.

Previous models have relaxed some of our simplifying assumptions, allowing a fixed (mating rate (λ)-independent) cost of distortion, and allowing the female mating rate (λ) to change as the distorter spreads[1,4,10,13,18-23]. We have explored these factors and found that our general conclusions: (i) are not altered; and (ii) do not depend on the distorter ($y_1$) allele spreading all the way to fixation (Scott, unpublished).

#### (2) Spread of an autosomal suppressor

We assume that the sex ratio distorter can be suppressed by an autosomal allele (suppressor), as has been found in many *Drosophila* species[24-27]. We base our model upon the biology of *Nmy*, which suppresses the X chromosome distorter *Dox* in *Drosophila simulans*. *Nmy* works by RNAi-mediated destruction of the distorter's mRNA transcripts. *Nmy* is dominant, and only expressed in the presence of the distorter (*Dox*)[28-30].

We denote the autosomal suppressor allele as *sup*, and the wild type non-suppressor allele as "+", after Vaz and Carvalho (2004), and for consistency across models. We assume that the suppressor (*sup*) is dominant, meaning individuals bearing at least one suppressor (*sup*) allele suffer no sperm death and consequentially no fertility loss or sex ratio distortion. We assume that the suppressor (*sup*) is only expressed in the presence of an active distorter (in $y_1/Y$ males). When the suppressor is expressed it leads to a cost, which reduces the probability ($V$) that an individual survives from zygote to adult[31-34], from $V_{normal}$=1 in individuals without an active suppressor, to $V_{suppression}$=1-$c_{sup}$ in individuals with one. The cost of suppression is a fixed cost ($c_{sup}$) of activating an RNAi pathway.

Assuming alternatively that the suppression cost affects fertility rather than viability does not qualitatively change our results (Scott, unpublished).

We ask when an autosomal suppressor (*sup*) will spread from rarity, given that an X chromosome distorter ($y_1$) is at fixation. Given that the suppressor only has phenotypic effects in $y_1/Y$ males, it will spread from rarity if $y_1/Y$ males bearing a suppressor (*sup*) have more mated offspring than $y_1/Y$ males lacking a suppressor (+/+). Assuming that the distorter and non-suppressor alleles are at fixation, and random mating, $y_1/Y$ males with a suppressor will have $V_{suppression}*F_{normal}*(½)*((1+k)/2)$ mated female offspring, and $V_{suppression}*F_{normal}*(½)*((1-k)/2)$ mated male offspring, leading to a total of $V_{suppression}*F_{normal}*(¼)$ mated offspring. $y_1/Y$ males lacking a suppressor will have a total of $2*V_{normal}*F_{drive}*((1-k)/2)*((1+k)/2)$ mated offspring. Suppressed $y_1$/Y males will therefore have more offspring, and the suppressor allele (*sup*) will spread from rarity, when the following condition is satisfied:

$$(F_{normal}/F_{drive})*(1/(1-k^2)) > (V_{normal}/V_{suppression}). \qquad \text{(S3)}$$

The overall cost of letting the distorter ($y_1$) go unsuppressed is a product of the costs to fertility ($F_{normal}/F_{drive}$) and offspring mating success ($1/(1-k^2)$). For a suppressor to spread, this must be greater than the viability cost of suppression ($V_{normal}/V_{suppression}$). Consequently, analogous to previous results, the suppressor (*sup*) will only spread when the distorter ($y_1$) leads to appreciable trait distortion[32,35-38].

A previous model asked whether female-biased sex ratio distortion can select for compensatory evolution on autosomes, such that the autosomes evolve to encode a male-biased sex ratio in the absence of the distorter[39]. It found that compensatory evolution does not evolve when the female-biased sex ratio distorter is transmitted into female offspring with 100% certainty, as is the case for X drivers acting in males. This is why we did not allow compensatory strategies to evolve on autosomes in our model, and only allowed autosomes to suppress the distorter.

#### (3) Consequences for organism trait values

We turn to the question of how distorter-suppressor dynamics affect sex ratio. When both the distorter ($y_1$) and suppressor (*sup*) are in a population, the genotypes they are in matters (epistasis), and so we explicitly track the frequencies of all 15 possible genotypes, with 15 recursions. The 15 equations represent the generational changes in each of the 15 possible genotypes. We let $p_{fi}$ and $q_{mi}$ be the proportion of the ith female genotype and the ith male genotype, respectively, in the current generation (Table S1). We let $p_{fi}$' and $q_{mi}$' be the frequencies of female and male genotypes in the next generation. The population sex ratio is given by the population proportion of females, $\sum p_f$. The equations are listed in Table S2. We note that, in the absence of the distorter ($y_1$), population sex ratio evolves to 0.5, and after this, genotype frequencies remain constant over time (Hardy-Weinberg equilibrium).

| | | **Females** | | | **Males** | |
|---|---|---|---|---|---|---|
| | | $y_0/y_0$ | $y_0/y_1$ | $y_1/y_1$ | $y_0$/Y | $y_1$/Y |
| ***sup / sup*** | ***Proportion*** | $p_{f1}$ | $p_{f4}$ | $p_{f7}$ | $p_{m1}$ | $p_{m4}$ |
| | ***Fertility, F*** | / | / | / | $F_{normal}$ | $F_{normal}$ |
| | ***Viability, V*** | 1 | 1 | 1 | 1 | $1-c_{sup}$ |

| | | | | | | |
|---|---|---|---|---|---|---|
| | ***Drive*** | / | / | / | 0.5 | 0.5 |
| ***sup / +*** | ***Proportion*** | $p_{f2}$ | $p_{f5}$ | $p_{f8}$ | $p_{m2}$ | $p_{m5}$ |
| | ***Fertility, F*** | / | / | / | $F_{normal}$ | $F_{normal}$ |
| | ***Viability, V*** | 1 | 1 | 1 | 1 | $1-c_{sup}$ |
| | ***Drive*** | / | / | / | 0.5 | 0.5 |
| ***+ / +*** | ***Proportion*** | $p_{f3}$ | $p_{f6}$ | $p_{f9}$ | $p_{m3}$ | $p_{m6}$ |
| | ***Fertility, F*** | / | / | / | $F_{normal}$ | $F_{drive}$ |
| | ***Viability, V*** | 1 | 1 | 1 | 1 | 1 |
| | ***Drive*** | / | / | / | 0.5 | $(1+k)/2$ |

**Table S1: Selection coefficients, drive values, and genotype frequency notation.** For each male and female genotype, its proportion in the population at generation *t*, and its probability of maturing from a zygote to an adult (viability, *V*) is given. For each male genotype, the proportion of X chromosomes in its sperm store (drive), and its probability of successfully fertilising the female's egg cell after copulation (fertility, *F*), is given. Male fertility (*F*) depends on the number of mates each female has per generation (λ), and is written in full in Equation S1. $k$ gives the proportion of a male's Y bearing sperm that are killed, and $c_{sup}$ gives the viability cost of distorter suppression.

| | |
|---|---|
| $T\ p_{f1}' =$ | $(p_{f1} + 0.5\ p_{f2} + 0.5\ p_{f4} + 0.25\ p_{f5})\ (0.5\ p_{m1} + 0.25\ p_{m2})\ F_{normal}$ |
| $T\ p_{f2}' =$ | $((0.5\ p_{f2} + p_{f3} + 0.25\ p_{f5} + 0.5\ p_{f6})\ (0.5\ p_{m1} + 0.25 p_{m2}) + (p_{f1} + 0.5\ p_{f2} + 0.5\ p_{f4} + 0.25\ p_{f5})\ (0.25\ p_{m2} + 0.5\ p_{m3}))\ F_{normal}$ |
| $T\ p_{f3}' =$ | $(0.5\ p_{f2} + p_{f3} + 0.25\ p_{f5} + 0.5\ p_{f6})\ (0.25 p_{m2} + 0.5\ p_{m3})\ F_{normal}$ |
| $T\ p_{f4}' =$ | $(0.5\ p_{f4} + 0.25\ p_{f5} + p_{f7} + 0.5\ p_{f8})\ (0.5\ p_{m1} + 0.25\ p_{m2}) + (p_{f1} + 0.5\ p_{f2} + 0.5\ p_{f4} + 0.25\ p_{f5})\ (p_{m4}/2 + p_{m5}/4)\ F_{normal}$ |
| $T\ p_{f5}' =$ | $((0.25\ p_{f5} + 0.5\ p_{f6} + 0.5\ p_{f8} + p_{f9})\ (0.5\ p_{m1} + 0.25\ p_{m2}) + (0.5\ p_{f4} + 0.25\ p_{f5} + p_{f7} + 0.5\ p_{f8})\ (0.25\ p_{m2} + 0.5\ p_{m3}))\ F_{normal} + (p_{f1} + 0.5\ p_{f2} + 0.5\ p_{f4} + 0.25\ p_{f5})\ (1/2\ (1 + k)\ F_{drive}\ p_{m6} + 1/4\ p_{m5}\ F_{normal}) + (0.5\ p_{f2} + p_{f3} + 0.25\ p_{f5} + 0.5\ p_{f6})\ (p_{m4}/2 + p_{m5}/4)\ F_{normal}$ |
| $T\ p_{f6}' =$ | $(0.25\ p_{f5} + 0.5\ p_{f6} + 0.5\ p_{f8} + p_{f9})\ (0.25\ p_{m2} + 0.5\ p_{m3})\ F_{normal} + (0.5\ p_{f2} + p_{f3} + 0.25\ p_{f5} + 0.5\ p_{f6})\ (1/2\ (1 + k)\ F_{drive}\ p_{m6} + 1/4\ p_{m5}\ F_{normal})$ |
| $T\ p_{f7}' =$ | $(0.5\ p_{f4} + 0.25\ p_{f5} + p_{f7} + 0.5\ p_{f8})\ (p_{m4}/2 + p_{m5}/4)\ F_{normal}$ |
| $T\ p_{f8}' =$ | $((0.5\ p_{f4} + 0.25\ p_{f5} + p_{f7} + 0.5\ p_{f8})\ (1/2\ (1 + k)\ F_{drive}\ p_{m6} + 1/4\ p_{m5}\ F_{normal}) + (0.25\ p_{f5} + 0.5\ p_{f6} + 0.5\ p_{f8} + p_{f9})\ (p_{m4}/2 + p_{m5}/4)\ F_{normal})$ |
| $T\ p_{f9}' =$ | $(0.25\ p_{f5} + 0.5\ p_{f6} + 0.5\ p_{f8} + p_{f9})\ (1/2\ (1 + k)\ F_{drive}\ p_{m6} + 1/4\ p_{m5} F_{normal})$ |
| $T\ p_{m1}' =$ | $(p_{f1} + 0.5\ p_{f2} + 0.5\ p_{f4} + 0.25\ p_{f5})\ ((0.5\ p_{m1} + 0.25\ p_{m2})\ F_{normal} + 1/2\ p_{m4}\ F_{normal} + 1/4\ p_{m5}\ F_{normal})$ |
| $T\ p_{m2}' =$ | $(p_{f1} + 0.5\ p_{f2} + 0.5\ p_{f4} + 0.25\ p_{f5})\ ((0.25\ p_{m2} + 0.5\ p_{m3}) F_{normal} + 1/2\ (1 - k)\ F_{drive}\ p_{m6} + 1/4\ p_{m5}\ F_{normal}) + (0.5\ p_{f2} + p_{f3} + 0.25\ p_{f5} + 0.5\ p_{f6})\ ((0.5\ p_{m1} + 0.25\ p_{m2}) F_{normal} + 1/2\ p_{m4}\ F_{normal} + 1/4\ p_{m5}\ F_{normal})$ |
| $T\ p_{m3}' =$ | $(0.5\ p_{f2} + p_{f3} + 0.25\ p_{f5} + 0.5\ p_{f6})\ ((0.25\ p_{m2} + 0.5\ p_{m3}) F_{normal} + 1/2\ (1 - k)\ F_{drive}\ p_{m6} + 1/4\ p_{m5}\ F_{normal})$ |

| | |
|---|---|
| T $p_{m4}'$ = | $V_{suppression}$ (0.5 $p_{f4}$ + 0.25 $p_{f5}$ + $p_{f7}$ + 0.5 $p_{f8}$) ((0.5 $p_{m1}$ + 0.25 $p_{m2}$)$F_{normal}$ + 1/2 $p_{m4}$ $F_{normal}$ + 1/4 $p_{m5}$ $F_{normal}$) |
| T $p_{m5}'$ = | $V_{suppression}$ ((0.5 $p_{f4}$ + 0.25 $p_{f5}$ + $p_{f7}$ + 0.5 $p_{f8}$) (0.25$F_{normal}$ $p_{m2}$ + 0.5$F_{normal}$ $p_{m3}$ + 1/2 (1- $k$) $F_{drive}$ $p_{m6}$ + 1/4 $p_{m5}$ $F_{normal}$) + (0.25 $p_{f5}$ + 0.5 $p_{f6}$ + 0.5 $p_{f8}$ + $p_{f9}$) (0.5 $p_{m1}$ + 0.25 $p_{m2}$ + 1/2 $p_{m4}$ $F_{normal}$ + 1/4 $p_{m5}$ $F_{normal}$)) |
| T $p_{m6}'$ = | (0.25 $p_{f5}$ + 0.5 $p_{f6}$ + 0.5 $p_{f8}$ + $p_{f9}$) ((0.25 $p_{m2}$ + 0.5 $p_{m3}$)$F_{normal}$ + 1/2 (1 – $k$) $F_{drive}$ $p_{m6}$ + 1/4 $p_{m5}$ $F_{normal}$). |

**Table S2: Recursions detailing the change in proportion of each genotype across one generation ($y_0$ and $y_1$ segregating at trait locus).** Notation is defined in Table S1. T is the sum of the right sides of the system of equations such that ∑p=1. It normalises the recursions to ensure that gene frequency changes reflect proportions.

To illustrate the logic of the equations, we derive one recursion explicitly. We derive the recursion for $p_{m5}'$, which gives the frequency, in the next generation, of males bearing the distorter (*$y_1$/Y*) and one suppressor allele (*sup/+*). We denote the current generation as G1 and the next generation as G2. A mating between *$y_1$/Y, +/+* males (at frequency $p_{m6}$) and *$y_1$/$y_1$, +/sup* females (at frequency $p_{f8}$) can give rise to individuals in G2 with our focal genotype *$y_1$/Y*, *sup/+*. Of all matings between males and females in G1 ($\sum_{i=1}^{6}\sum_{j=1}^{9} p_{mi} * p_{fj}$), these matings occur in the proportion $(p_{m6} * p_{f8})/(\sum_{i=1}^{6}\sum_{j=1}^{9} p_{mi} * p_{fj})$ of cases. Copulation success, in which the egg is successfully fertilised to form a zygote, depends on the fertility of the male, which in our case is $F_{drive}$ (Equation S1a). Of all the zygotes produced by the population in G1, our parents will contribute the proportion $(p_{m6} * p_{f8})/(\sum_{i=1}^{6}\sum_{j=1}^{9} p_{mi} * p_{fj})$) * ($F_{drive}$ / $\sum_{i=1}^{6} p_{mi} F_{mi}$) of them, where $\sum_{i=1}^{6} p_{mi} F_{mi}$ gives average male fertility. These zygotes will have the focal offspring genotype (*$y_1$/Y, sup/+*) if they inherit a Y+ gamete from the father (with probability (1-$k$)/2) and a *$y_1$sup* gamete from the mother (with probability 1/2), meaning the proportion of zygotes in the population with the focal genotype, (*$y_1$/Y, sup/+*), stemming from copulations between our focal parents, is $(p_{m6} * p_{f8})/(\sum_{i=1}^{6}\sum_{j=1}^{9} p_{mi} * p_{fj})$) * ($F_{drive}$ / $\sum_{i=1}^{6} p_{mi}\, F_{mi}$) * (1+$k$)/4. Finally, only the proportion $V_{suppression}$ of these focal zygotes (*$y_1$/Y, sup/+*) will successfully mature to adulthood in G2, meaning the proportion of mature adults in G2 that have the focal genotype (*$y_1$/Y, sup/+*) and arose from copulations between *$y_1$/Y, +/+* males and *$y_1$/$y_1$, +/sup* females in G1 is given by

$(p_{m6} * p_{f8})/(\sum_{i=1}^{6}\sum_{j=1}^{9} p_{mi} * p_{fj})$) * ($F_{drive}$ / $\sum_{i=1}^{6} p_{mi}\, F_{mi}$) * (1+$k$)/4 * ($V_{suppression}$ / $\sum p * V$), where $\sum p * V$ is the average viability of individuals. We simplify the expression by making the substitution $T = (\sum_{i=1}^{6}\sum_{j=1}^{9} p_{mi} * p_{fj})(\sum_{i=1}^{6} p_{mi}\, F_{mi})(\sum p * V)$, our normalisation factor. We now need to sum over all possible parental copulations that can give rise to *$y_1$/Y, +/sup* offspring. Doing so gives the frequency of the *$y_1$/Y, +/sup* genotype in the next generation, $p_{m5}'$, written in full in Table S2.

We iterated these recursions to find the distorter (*$y_1$*) and suppressor (*sup*) frequencies, and the population sex ratio ($\sum p_f$), at equilibrium (Fig. S3). When we introduced both the distorter and suppressor at low frequencies, we confirmed our above results that the distorter (*$y_1$*) initially spreads to fixation, and that the suppressor allele (*sup*) only invades and reaches high frequencies if it is suppressing a strong distorter (high $k$).

We used our recursions to examine whether the spread of the suppressor led to the subsequent loss of the distorter. As the suppressor increases in frequency, the population sex ratio becomes less biased, and the fitness benefit of further distorter suppression is reduced (negative frequency dependence). This means that, when females are singly mated (λ=1), the rise of the suppressor allele towards some

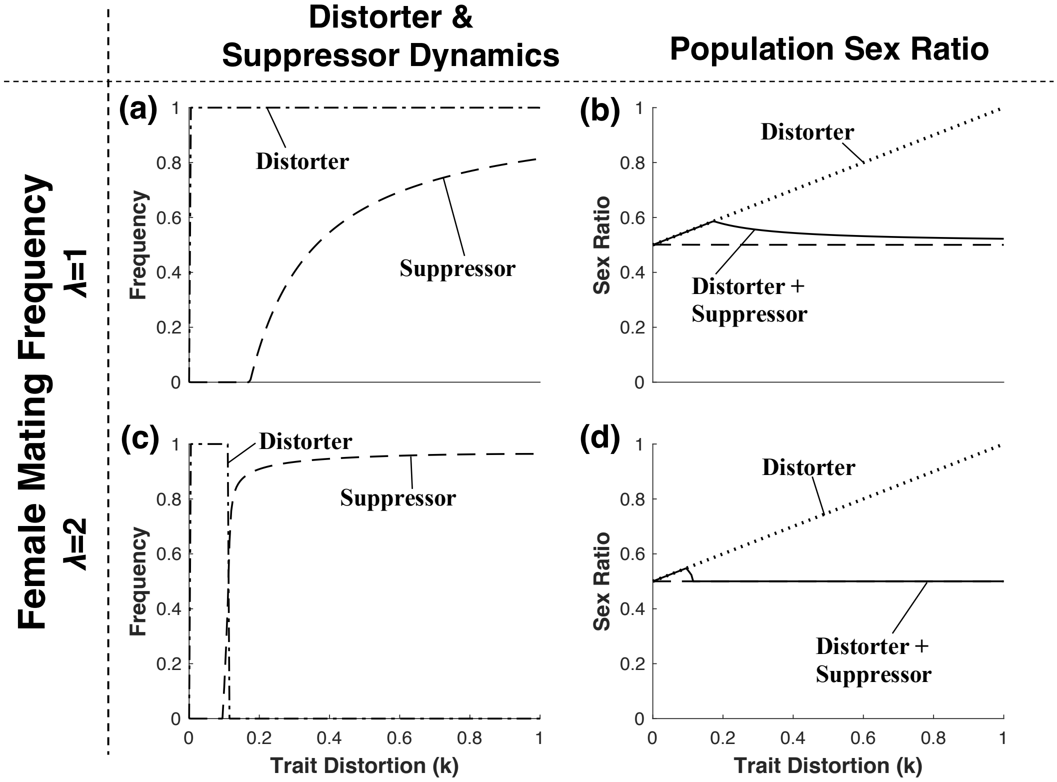


**Figure S3. Drive-suppressor coevolution and resulting sex ratio.** A sex ratio distorter ($y_1$) and its suppressor (*sup*) are introduced from rarity. The left-hand column shows the resulting equilibrium distorter ($y_1$; *dot-dash* line) and suppressor (*sup*; dashed line) frequencies, for different distorter strengths ($0<k\leq1$). The right-hand column shows the resulting population average sex ratio. The dotted and dashed lines are plotted for reference, to show, respectively, the sex ratio $(1+k)/2$) that arises in the absence of suppression, and the sex ratio (½) that arises in the absence of the distorter ($y_1$). The top row shows results for when females are singly mated (λ=1) and the bottom row shows results for when females are doubly mated (λ=2). The numerical solutions assume that the cost of suppression is $c_{sup}$=0.03. We see that equilibrium suppressor (*sup*) frequency is greater for strong (higher-$k$) drivers, resulting in full ($\lambda$=2) or partial ($\lambda$=1) restoration of the individual optimal sex ratio (0.5) for strong (higher-$k$) drivers.

nonzero equilibrium frequency does not cause subsequent loss of the distorter ($y_1$), which remains at fixation (Fig. S3a). When females are multiply mated (λ>1), there is an additional fertility cost of distortion, and so the suppressor continues to spread, to a higher equilibrium frequency, until the distorter ($y_1$) is lost completely from the population (Fig. S3c).

We also considered the overall consequences of the distorter-suppressor dynamics for the sex ratio. For weak distorters (low $k$), suppressors do not spread. Consequently, there is sex ratio distortion, but it is negligible. For distorters of intermediate strength (intermediate $k$, e.g. $k \approx 0.2$), suppressors are still at low population frequency, and so there can be greater sex ratio distortion. For strong distorters (high $k$), suppressors spread to high population frequency, and so there is little (when $\lambda=1$) or no (when $\lambda>1$) sex ratio distortion. Consequently, the extent that the sex ratio deviates from the individual optimum of equal investment in the sexes: (a) shows a domed relationship with the extent of distortion ($k$); and (b) will often be negligible[37] (Fig. S3b & Fig. S3d). It should be noted that, in reality, the population is a mixture of two types of individual, one adopting a sex ratio of ½ and the other adopting a distorted sex ratio of $(1+k)/2$, and here we are capturing the population average deviation of individuals from the optimal sex ratio.

The case of singly mated females ($\lambda=1$) is of special interest because there is no fertility cost of trait distortion ($F_{normal}=F_{drive}$), meaning the individual level cost of bearing the selfish genetic element ($y_1$) arises *solely* because an individual level trait (sex ratio) is suboptimal (not ½). Sex ratio distortion is often negligible even in this special case ($\lambda=1$), indicating that the parliament of genes can act for the sole purpose of trait (sex ratio) restoration, without the additional incentive of fertility recovery.

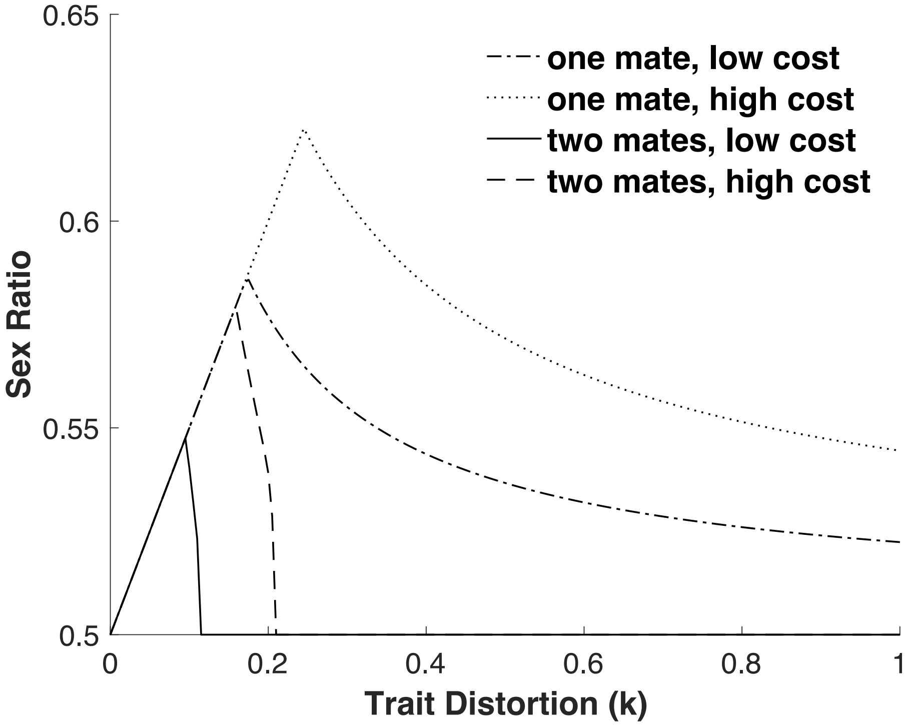


**Figure S4. Mating frequency and cost of suppression.** A sex ratio distorter ($y_1$) and its suppressor (*sup*) are introduced from rarity. Equilibrium sex ratio is plotted, across multiple trails where $y_1$ has different levels of drive ($0<k\leq1$), and where different assumptions are made about the cost of suppression ($c_{sup}$) and female mating rate (λ). The four parameter regimes plotted assume single (λ=1) or double (λ=2) female mating; and, low ($c_{sup}$=0.03) or high ($c_{sup}$=0.06) viability cost of distorter suppression. The sex ratio is more easily distorted when the suppressor is costlier ($c_{sup}$) and when females mating rate is lower (λ).

Additionally, we considered the effects of model parameters on sex ratio. We found that increasing the rate of female mating (λ) and decreasing the cost of suppression ($c_{sup}$) both led to a reduced tolerance of drive, and a correspondingly reduced level of sex ratio distortion (Fig. S4).

Finally, we considered how far sex ratio can be distorted in the time period after the distorter initially invades and before the distorter is suppressed and purged from the

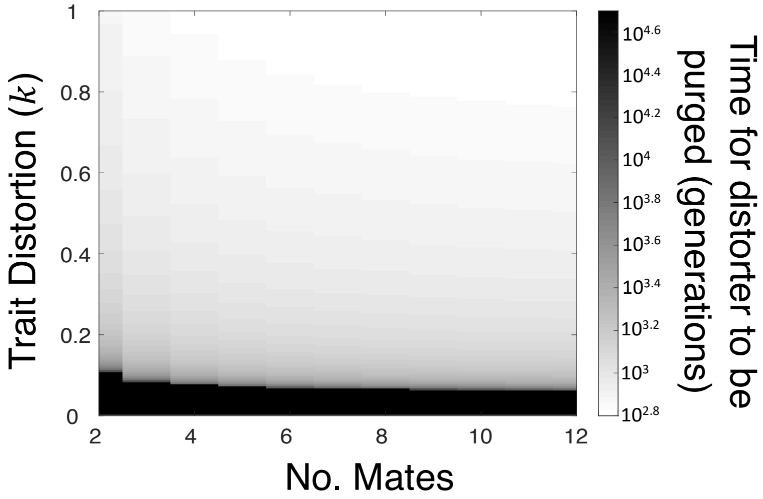


**Figure S5. Non-equilibrium sex ratio distortion.** An X driver ($y_1$) is introduced from rarity alongside its suppressor (*sup*). The proportion of Y-bearing sperm killed by the X driver ($k$) is varied alongside number of times each female mates per generation ($\lambda$). The cost of suppression is fixed at $c_{sup}$=0.03. We consider distorters that are purged from the population, after being suppressed, at equilibrium ($\lambda$>1). We plot the number of generations (on a $\log_{10}$ scale) until equilibrium is reached (trials that did not equilibrate by 50,000 generations were capped). We see that stronger distorters (high $k$) are purged at a faster rate, reducing the potential for non-equilibrium sex ratio distortion. Increased female mating (high $\lambda$) increases the fertility cost of distortion, meaning distorters are purged at a faster rate.

population. We iterated our recursions and timed how many generations it took to reach equilibrium. We found that stronger distorters (higher $k$) are suppressed and purged from the population more quickly, especially at higher female mating rates ($\lambda$) where the fertility cost of sex ratio distortion is greater (Fig. S5).

#### 4) Evolution of trait distortion

In the above analyses, we assumed that the strength of the distorter ($y_1$) was a fixed parameter ($k$). We now consider the consequence of allowing the level of distortion to evolve[16]. We first consider the scenario in which there is no suppressor. We take a game theoretical approach to find the evolutionarily stable level of X chromosome distortion ($k^*$) in the absence of suppression. We assume a population where all males have an X chromosome with the same level of distortion, denoted by a capital *K.* We then assume that a mutation arises in the X chromosome of one male in the population that causes it to assume a new level of distortion, denoted by $\breve{k}$. We wish to find the level of distortion that, when adopted by every X chromosome in the population, cannot be invaded by the mutant X chromosome adopting a different level of distortion. This level of distortion ($k^*$) represents the evolutionarily stable strategy (ESS)[40].

Distorters have no effect in females, so the fitness of the mutant distorter depends only on its action in males. The male bearing the mutant distorter has fertility given by its proportional sperm contribution to a female mate's sperm store: $\left(\frac{\left(1-{\breve{k}}/{2}\right)}{\left(1-{\breve{k}}/{2}\right)+(\lambda-1)\left(1-{K}/{2}\right)}\right)$. The mutant distorter is passed into (1+$\breve{k}$)/2 offspring, and so the fitness of the mutant X chromosome is proportional to: $w = \frac{(1+\breve{k})}{2}\left(\frac{\left(1-{\breve{k}}/{2}\right)}{\left(1-{\breve{k}}/{2}\right)+(\lambda-1)\left(1-{K}/{2}\right)}\right)$. The ESS level of X chromosome distortion is the value of $k^*$ that satisfies $\frac{dw}{d\breve{k}} = 0$ and $\frac{d^2w}{d\breve{k}^2} < 0$ when $\breve{k}$=$K$=$k^*$, and is given by:

$$k^*=(\lambda+1)/(2\lambda-1). \qquad \text{(S4)}$$

When females mate singly (λ=1) or doubly (λ=2), maximal distortion is favoured ($k^*$=1), resulting in population collapse due to lack of males. As female mating frequency increases to λ≥3, the increased fertility cost of distortion means that the

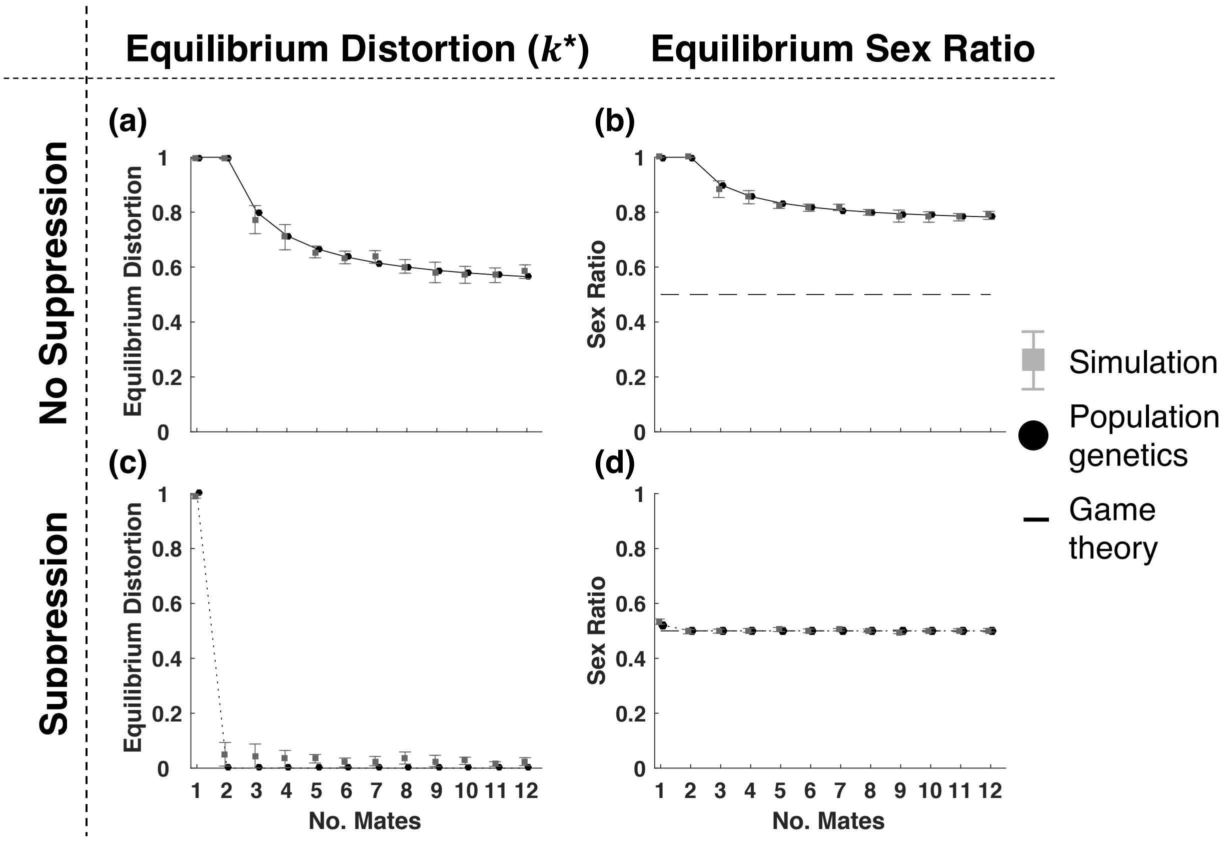


**Figure S6. Equilibrium distorter strength and sex ratio.** (a) In the absence of a suppressor (*sup*), equilibrium distortion decreases with the number of mates each female has per generation ($\lambda$). (b) The distortion at equilibrium causes a more significantly distorted equilibrium sex ratio at lower female mating rates. A dashed line shows the sex ratio that would evolve in the absence of selfish genes (½). (c) In the presence of suppression, maximally distorting (but largely suppressed) distorters ($k^*$=1) evolve when females are singly mated ($\lambda$=1); otherwise, non-distorters ($k^*$=0) evolve. (d) Owing to the spread of suppressors, sex ratio is completely ($\lambda$>1) or partially ($\lambda$=1) recovered at equilibrium. (c) and (d) assumed a small cost of suppression ($c_{\mathrm{sup}}$=0.03). For all graphs, the results of the simulation (grey boxes) are plotted alongside the population genetics result (black circles). For (a) and (b), the result of a game theoretic analysis is also plotted (solid line). All methods give the same equilibrium level of distortion and sex ratio. The error bars show one standard deviation from the mean over 10 trials of the simulation.

equilibrium level of distortion ($k^*$) decreases[16], until it plateaus at the minimum of $k^*$=0.5 as $\lambda \to \infty$ (Fig. S6a). The game theoretic equilibrium is verified in fully dynamical population genetic and agent-based simulation models, as described below (Fig. S6a).

We now consider what sex ratio will evolve in the presence of a suppressor (*sup*). We assume that a mutant X chromosome distorter ($y_2$) arises from a mutation on the old sex ratio distorter ($y_1$), and kills a different proportion of sperm when unsuppressed ($\hat{k} \neq k$), biasing individual sex ratio by $(1+\hat{k})/2$ and reducing ejaculate size to $1-\hat{k}/2$. We assume that $y_2$ and $y_1$ share a similar genetic and mechanistic basis of drive, such that the mutant distorting X chromosome ($y_2$) is suppressed by the same suppressor allele (*sup*)[41-44]. In Table S4, we display 27 recursions to describe the generational changes in genotype frequencies when the alleles $y_1$, $y_0$, $y_2$, $Y$, + and *sup* are segregating in a population (notation defined in Table S1 & S3). We note that, in the absence of the distorters ($y_1$ and $y_2$), population sex ratio evolves to 0.5, and after this, genotype frequencies remain constant over time (Hardy-Weinberg equilibrium). These equations reduce to those in Table S2 when genotypes bearing $y_2$ are set to zero.

| | | **Females** | | **Males** | |
|---|---|---|---|---|---|
| | | $y_0/y_2$ | $y_1/y_2$ | $y_2/y_2$ | $y_2/Y$ |
| ***sup / sup*** | ***Proportion*** | $p_{f10}$ | $p_{f13}$ | $p_{f16}$ | $p_{m7}$ |
| | ***Fertility, F*** | / | / | / | $F_{normal}$ |
| | ***Viability, V*** | 1 | 1 | 1 | $1-c_{sup}$ |
| | ***Drive*** | / | / | / | 0.5 |
| ***sup / +*** | ***Proportion*** | $p_{f11}$ | $p_{f14}$ | $p_{f17}$ | $p_{m8}$ |
| | ***Fertility, F*** | / | / | / | $F_{normal}$ |
| | ***Viability, V*** | 1 | 1 | 1 | $1-c_{sup}$ |
| | ***Drive*** | / | / | / | 0.5 |

| | | | | | |
|---|---|---|---|---|---|
| +/+ | ***Proportion*** | $p_{f12}$ | $P_{f15}$ | $p_{f18}$ | $p_{m9}$ |
| | ***Fertility, F*** | / | / | / | $F_{driveMut}$ |
| | ***Viability, V*** | 1 | 1 | 1 | 1 |
| | ***Drive*** | / | / | / | $(1+\hat{k})/2$ |

**Table S3: Further Selection coefficients, drive values, and genotype frequency notation.** For each male and female genotype, its proportion in the population at generation *t*, and its probability of maturing from a zygote to an adult (viability, *V*) is given. For each male genotype, the proportion of X chromosomes in its sperm store (drive), and its probability of successfully fertilising the female's egg cell after copulation (fertility, *F*), is given. Male fertility (*F*) depends on the number of mates each female has per generation (λ) according to:

$$F_{normal} = \sum_{i=0}^{\lambda-1} \sum_{j=0}^{\lambda-i-1} \frac{1}{1+\left(1-\frac{k}{2}\right)i+j+(\lambda-i-j-1)(1-\frac{\hat{k}}{2})} l^i (1-l-n)^j n^{\lambda-i-j-1},$$

$$F_{drive} = \sum_{i=0}^{\lambda-1} \sum_{j=0}^{\lambda-i-1} \frac{1-\frac{k}{2}}{\left(1-\frac{k}{2}\right)(i+1)+j+(\lambda-i-j-1)(1-\frac{\hat{k}}{2})} l(t)^i (1-l-n)^j n^{\lambda-i-j-1},$$

$$F_{driveMut} = \sum_{i=0}^{\lambda-1} \sum_{j=0}^{\lambda-i-1} \frac{1-\frac{\hat{k}}{2}}{\left(1-\frac{k}{2}\right)i+j+(\lambda-i-j)(1-\frac{\hat{k}}{2})} l^i (1-l-n)^j n^{\lambda-i-j-1}.$$

*l*, *n*, and *1-l-n*, are, respectively, the proportions of males in the population with: an unsuppressed $y_1$; an unsuppressed $y_2$; neither of these (all other males). $k$ and $\hat{k}$ respectively give the proportion of a male's Y bearing sperm that are killed by an unsuppressed $y_1$ and $y_2$ distorter, and $c_{sup}$ gives the viability cost of distorter suppression.

| | |
|---|---|
| T $p_{f1}'$ = | $(p_{f1} + 0.5\ p_{f10} + 0.25\ p_{f11} + 0.5\ p_{f2} + 0.5\ p_{f4} + 0.25\ p_{f5})\ (0.5\ p_{m1} + 0.25\ p_{m2})\ F_{normal}$ |
| T $p_{f2}'$ = | $((0.25\ p_{f11} + 0.5\ p_{f12} + 0.5\ p_{f2} + p_{f3} + 0.25\ p_{f5} + 0.5\ p_{f6})\ (0.5\ p_{m1} + 0.25\ p_{m2})$ + $(p_{f1} + 0.5\ p_{f10} + 0.25\ p_{f11} + 0.5\ p_{f2} + 0.5\ p_{f4} + 0.25\ p_{f5})\ (0.25\ p_{m2} + 0.5\ p_{m3}))\ F_{normal}$ |
| T $p_{f3}'$ = | $(0.25\ p_{f11} + 0.5\ p_{f12} + 0.5\ p_{f2} + p_{f3} + 0.25\ p_{f5} + 0.5\ p_{f6})\ (0.25\ p_{m2} + 0.5\ p_{m3})\ F_{normal}$ |
| T $p_{f4}'$ = | $(0.5\ p_{f13} + 0.25\ p_{f14} + 0.5\ p_{f4} + 0.25\ p_{f5} + p_{f7} + 0.5\ p_{f8})\ (0.5\ p_{m1} + 0.25\ p_{m2})\ F_{normal}$ + 1/4 $(p_{f1} + 0.5\ p_{f10} + 0.25\ p_{f11} + 0.5\ p_{f2} + 0.5\ p_{f4} + 0.25\ p_{f5})\ (2\ p_{m4}\ F_{drive} + p_{m5}\ F_{normal})$ |
| T $p_{f5}'$ = | $(0.25\ p_{f14} + 0.5\ p_{f15} + 0.25\ p_{f5} + 0.5\ p_{f6} + 0.5\ p_{f8} + p_{f9})\ (0.5\ p_{m1} + 0.25\ p_{m2})\ F_{normal}$ + $(0.5\ p_{f13} + 0.25\ p_{f14} + 0.5\ p_{f4} + 0.25\ p_{f5} + p_{f7} + 0.5\ p_{f8})\ (0.25\ p_{m2} + 0.5\ p_{m3})\ F_{normal}$ + 1/4 $(0.25\ p_{f11} + 0.5\ p_{f12} + 0.5\ p_{f2} + p_{f3} + 0.25\ p_{f5} + 0.5\ p_{f6})\ (2\ p_{m4}\ F_{drive} + p_{m5}\ F_{drive})$ + 1/4 $(p_{f1} + 0.5\ p_{f10} + 0.25\ p_{f11} + 0.5\ p_{f2} + 0.5\ p_{f4} + 0.25\ p_{f5})\ (-2\ (1 + k)\ F_{drive}\ p_{m6} + p_{m5}\ F_{normal})$ |
| T $p_{f6}'$ = | $(0.25\ p_{f14} + 0.5\ p_{f15} + 0.25\ p_{f5} + 0.5\ p_{f6} + 0.5\ p_{f8} + p_{f9})\ (0.25\ p_{m2} + 0.5\ p_{m3})\ F_{normal}$ + 1/4 $(0.25\ p_{f11} + 0.5\ p_{f12} + 0.5\ p_{f2} + p_{f3} + 0.25\ p_{f5} + 0.5\ p_{f6})\ (-2\ (1 + k)\ F_{drive}\ p_{m6} + p_{m5}\ F_{normal})$ |
| T $p_{f7}'$ = | 1/4 $(0.5\ p_{f13} + 0.25\ p_{f14} + 0.5\ p_{f4} + 0.25\ p_{f5} + p_{f7} + 0.5\ p_{f8})\ (2\ p_{m4}\ F_{drive} + p_{m5}\ F_{normal})$ |
| T $p_{f8}'$ = | 1/4 $((0.25\ p_{f14} + 0.5\ p_{f15} + 0.25\ p_{f5} + 0.5\ p_{f6} + 0.5\ p_{f8} + p_{f9})\ (2\ p_{m4}\ F_{drive} + p_{m5}\ F_{normal})$ + $(0.5\ p_{f13} + 0.25\ p_{f14} + 0.5\ p_{f4} + 0.25\ p_{f5} + p_{f7} + 0.5\ p_{f8})\ (-2\ (1 + k)\ F_{drive}\ p_{m6} + p_{m5}$ |

| | |
|---|---|
| | $F_{normal}))$ |
| $T\ p_{f9}' =$ | $1/4\ (0.25\ p_{f14} + 0.5\ p_{f15} + 0.25\ p_{f5} + 0.5\ p_{f6} + 0.5\ p_{f8} + p_{f9})\ (-2\ (1 + k)\ F_{drive}\ p_{m6} + p_{m5}\ F_{normal})$ |
| $T\ p_{f10}' =$ | $(0.5\ p_{f10} + 0.25\ p_{f11} + 0.5\ p_{f13} + 0.25\ p_{f14} + p_{f16} + 0.5\ p_{f17})\ (0.5\ p_{m1} + 0.25\ p_{m2})\ F_{normal} - 1/4\ (p_{f1} + 0.5\ p_{f10} + 0.25\ p_{f11} + 0.5\ p_{f2} + 0.5\ p_{f4} + 0.25\ p_{f5})\ (-2\ p_{m7} - p_{m8})\ F_{driveMut}$ |
| $T\ p_{f11}' =$ | $((0.25\ p_{f11} + 0.5\ p_{f12} + 0.25\ p_{f14} + 0.5\ p_{f15} + 0.5\ p_{f17} + p_{f18})\ (0.5\ p_{m1} + 0.25\ p_{m2}) + (0.5\ p_{f10} + 0.25\ p_{f11} + 0.5\ p_{f13} + 0.25\ p_{f14} + p_{f16} + 0.5\ p_{f17})\ (0.25\ p_{m2} + 0.5\ p_{m3}))\ F_{normal}$ |
| $T\ p_{f12}' =$ | $(0.25\ p_{f11} + 0.5\ p_{f12} + 0.25\ p_{f14} + 0.5\ p_{f15} + 0.5\ p_{f17} + p_{f18})\ (0.25\ p_{m2} + 0.5\ p_{m3})\ F_{normal} + 1/4\ (0.25\ p_{f11} + 0.5\ p_{f12} + 0.5\ p_{f2} + p_{f3} + 0.25\ p_{f5} + 0.5\ p_{f6})\ (-2\ (1 + \hat{k})\ F_{driveMut}\ p_{m9} + p_{m8}\ F_{driveMut})$ |
| $T\ p_{f13}' =$ | $1/4\ ((0.5\ p_{f10} + 0.25\ p_{f11} + 0.5\ p_{f13} + 0.25\ p_{f14} + p_{f16} + 0.5\ p_{f17})\ (2\ p_{m4}\ F_{drive} + p_{m5}\ F_{normal}) - (0.5\ p_{f13} + 0.25\ p_{f14} + 0.5\ p_{f4} + 0.25\ p_{f5} + p_{f7} + 0.5\ p_{f8})\ (-2\ p_{m7} - p_{m8})\ F_{driveMut})$ |
| $T\ p_{f14}' =$ | $1/4\ ((0.25\ p_{f11} + 0.5\ p_{f12} + 0.25\ p_{f14} + 0.5\ p_{f15} + 0.5\ p_{f17} + p_{f18})\ (2\ p_{m4}\ F_{drive} + p_{m5}\ F_{normal}) + (0.5\ p_{f10} + 0.25\ p_{f11} + 0.5\ p_{f13} + 0.25\ p_{f14} + p_{f16} + 0.5\ p_{f17})\ (-2\ (1 + k)\ F_{drive}\ p_{m6} + p_{m5}\ F_{normal}) + (0.5\ p_{f13} + 0.25\ p_{f14} + 0.5\ p_{f4} + 0.25\ p_{f5} + p_{f7} + 0.5\ p_{f8})\ (-2\ (1 + \hat{k})\ F_{driveMut}\ p_{m9} + p_{m8}\ F_{driveMut}) - (0.25\ p_{f14} + 0.5\ p_{f15} + 0.25\ p_{f5} + 0.5\ p_{f6} + 0.5\ p_{f8} + p_{f9})\ (-2\ p_{m7} - p_{m8})\ F_{driveMut})$ |
| $T\ p_{f15}' =$ | $1/4\ ((0.25\ p_{f11} + 0.5\ p_{f12} + 0.25\ p_{f14} + 0.5\ p_{f15} + 0.5\ p_{f17} + p_{f18})\ (-2\ (1 + k)\ F_{drive}\ p_{m6} + p_{m5}\ F_{normal}) + (0.25\ p_{f14} + 0.5\ p_{f15} + 0.25\ p_{f5} + 0.5\ p_{f6} + 0.5\ p_{f8} + p_{f9})\ (-2\ (1 + \hat{k})\ F_{driveMut}\ p_{m9} + p_{m8}\ F_{driveMut}))$ |
| $T\ p_{f16}' =$ | $1/4\ (0.5\ p_{f10} + 0.25\ p_{f11} + 0.5\ p_{f13} + 0.25\ p_{f14} + p_{f16} + 0.5\ p_{f17})\ (-2\ p_{m7} - p_{m8})\ F_{driveMut}$ |
| $T\ p_{f17}' =$ | $1/4\ ((0.5\ p_{f10} + 0.25\ p_{f11} + 0.5\ p_{f13} + 0.25\ p_{f14} + p_{f16} + 0.5\ p_{f17})\ (-2\ (1 + \hat{k})\ F_{driveMut}\ p_{m9} + p_{m8}\ F_{driveMut}) - (0.25\ p_{f11} + 0.5\ p_{f12} + 0.25\ p_{f14} + 0.5\ p_{f15} + 0.5\ p_{f17} + p_{f18})\ (-2\ p_{m7} - p_{m8})\ F_{driveMut})$ |
| $T\ p_{f18}' =$ | $1/4\ (0.25\ p_{f11} + 0.5\ p_{f12} + 0.25\ p_{f14} + 0.5\ p_{f15} + 0.5\ p_{f17} + p_{f18})\ (-2\ (1 + \hat{k})\ F_{driveMut}\ p_{m9} + p_{m8}\ F_{driveMut})$ |
| $T\ p_{m1}' =$ | $1/4\ (p_{f1} + 0.5\ p_{f10} + 0.25\ p_{f11} + 0.5\ p_{f2} + 0.5\ p_{f4} + 0.25\ p_{f5})\ ((2\ p_{m1} + p_{m2})\ F_{normal} - 2\ F_{drive}\ p_{m4} + p_{m5}\ F_{normal} + 2\ p_{m7}\ F_{driveMut} + p_{m8}\ F_{driveMut})$ |
| $T\ p_{m2}' =$ | $1/4\ ((p_{f1} + 0.5\ p_{f10} + 0.25\ p_{f11} + 0.5\ p_{f2} + 0.5\ p_{f4} + 0.25\ p_{f5})\ ((p_{m2} + 2\ p_{m3})\ F_{normal} + 2\ (-1 + k)\ F_{drive}\ p_{m6} + 2\ (-1 + \hat{k})\ F_{driveMut}\ p_{m9} + p_{m5}\ F_{normal} + p_{m8}\ F_{driveMut}) + (0.25\ p_{f11} + 0.5\ p_{f12} + 0.5\ p_{f2} + p_{f3} + 0.25\ p_{f5} + 0.5\ p_{f6})\ ((2\ p_{m1} + p_{m2})\ F_{normal} - 2\ F_{drive}\ p_{m4} + p_{m5}\ F_{normal} + 2\ p_{m7}\ F_{driveMut} + p_{m8}\ F_{driveMut}))$ |
| $T\ p_{m3}' =$ | $1/4\ (0.25\ p_{f11} + 0.5\ p_{f12} + 0.5\ p_{f2} + p_{f3} + 0.25\ p_{f5} + 0.5\ p_{f6})\ ((p_{m2} + 2\ p_{m3})\ F_{normal} + 2\ (-1 + k)\ F_{drive}\ p_{m6} + 2\ (-1 + \hat{k})\ F_{driveMut}\ p_{m9} + p_{m5}\ F_{normal} + p_{m8}\ F_{driveMut})$ |
| $T\ p_{m4}' =$ | $1/4\ V_{suppression}\ (0.5\ p_{f13} + 0.25\ p_{f14} + 0.5\ p_{f4} + 0.25\ p_{f5} + p_{f7} + 0.5\ p_{f8})\ ((2\ p_{m1} + p_{m2})\ F_{normal} - 2\ F_{drive}\ p_{m4} + p_{m5}\ F_{normal} + 2\ p_{m7}\ F_{driveMut} + p_{m8}\ F_{driveMut})$ |
| $T\ p_{m5}' =$ | $1/4\ V_{suppression}\ ((0.5\ p_{f13} + 0.25\ p_{f14} + 0.5\ p_{f4} + 0.25\ p_{f5} + p_{f7} + 0.5\ p_{f8})\ ((p_{m2} + 2\ p_{m3})\ F_{normal} + 2\ (-1 + k)\ F_{drive}\ p_{m6} + 2\ (-1 + \hat{k})\ F_{driveMut}\ p_{m9} + p_{m5}\ F_{normal} + p_{m8}\ F_{driveMut}) + (0.25\ p_{f14} + 0.5\ p_{f15} + 0.25\ p_{f5} + 0.5\ p_{f6} + 0.5\ p_{f8} + p_{f9})\ ((2\ p_{m1} + p_{m2})\ F_{normal} - 2\ F_{drive}\ p_{m4} + p_{m5}\ F_{normal} + 2\ p_{m7}\ F_{driveMut} + p_{m8}\ F_{driveMut}))$ |
| $T\ p_{m6}' =$ | $1/4\ (0.25\ p_{f14} + 0.5\ p_{f15} + 0.25\ p_{f5} + 0.5\ p_{f6} + 0.5\ p_{f8} + p_{f9})\ ((p_{m2} + 2\ p_{m3})\ F_{normal} + 2\ (-1 + k)\ F_{drive}\ p_{m6} + 2\ (-1 + \hat{k})\ F_{driveMut}\ p_{m9} + p_{m5}\ F_{normal} + p_{m8}\ F_{driveMut})$ |
| $T\ p_{m7}' =$ | $1/4\ V_{suppression}\ (0.5\ p_{f10} + 0.25\ p_{f11} + 0.5\ p_{f13} + 0.25\ p_{f14} + p_{f16} + 0.5\ p_{f17})\ ((2\ p_{m1} + p_{m2})\ F_{normal} - 2\ F_{drive}\ p_{m4} + p_{m5}\ F_{normal} + 2\ p_{m7}\ F_{driveMut} + p_{m8}\ F_{driveMut})$ |
| $T\ p_{m8}' =$ | $1/4\ V_{suppression}\ ((0.5\ p_{f10} + 0.25\ p_{f11} + 0.5\ p_{f13} + 0.25\ p_{f14} + p_{f16} + 0.5\ p_{f17})\ ((p_{m2} + 2\ p_{m3})\ F_{normal} + 2\ (-1 + k)\ F_{drive}\ p_{m6} + 2\ (-1 + \hat{k})\ F_{driveMut}\ p_{m9} + p_{m5}\ F_{normal} + p_{m8}\ F_{driveMut}) + (0.25\ p_{f11} + 0.5\ p_{f12} + 0.25\ p_{f14} + 0.5\ p_{f15} + 0.5\ p_{f17} + p_{f18})\ ((2\ p_{m1} + p_{m2})\ F_{normal} - 2\ F_{drive}\ p_{m4} + p_{m5}\ F_{normal} + 2\ p_{m7}\ F_{driveMut} + p_{m8}\ F_{driveMut}))$ |
| $T\ p_{m9}' =$ | $1/4\ (0.25\ p_{f11} + 0.5\ p_{f12} + 0.25\ p_{f14} + 0.5\ p_{f15} + 0.5\ p_{f17} + p_{f18})\ ((p_{m2} + 2\ p_{m3})\ F_{normal} + 2\ (-1 + k)\ F_{drive}\ p_{m6} + 2\ (-1 + \hat{k})\ F_{driveMut}\ p_{m9} + p_{m5}\ F_{normal} + p_{m8}\ F_{driveMut})$ |

**Table S4: Recursions detailing the change in proportion of each genotype across one generation ($y_0$, $y_1$ and $y_2$ segregating at trait locus).** Notation is defined in Table S1 & S3. T is the sum of the right sides of the system of equations such that ∑p=1. It normalises the recursions to ensure that gene frequency changes reflect proportions.

We work out the evolved level of sex ratio distortion, under the assumption that distortion is initially low, and the additional assumption of weak selection. We assume the mutant distorter ($y_2$) is only slightly stronger than the distorter from which it is derived ($y_1$), so that $\hat{k}=k+\delta$, where $\delta$ is positive and very small ($\delta$-*weak selection*[45]). We see if a mutant distorter can spread by iterating our recursions in Table S4 until equilibrium is reached. If the stronger distorter ($y_2$) displaces the weaker one ($y_1$), we introduce a further mutant distorter and iterate our equations again. We elucidate the equilibrium distorter strength ($k^*$) by successively introducing mutant distorters ($y_2$) until one fails to invade, at which point the equilibrium level of distortion ($k^*$) has been reached.

We find that, in the presence of the suppressor allele (*sup*), weakly distorting X chromosomes (low-$k$) can evade suppression and successfully distort sex ratio. These weak distorters will be displaced by slightly more distorting mutants ($y_2$). If the cost of suppression ($c_{sup}$) is sufficiently low, this displacement causes the frequency of the suppressor allele (*sup*) to increase in response. This trend means that, given sequential mutations on the X chromosome to increase sex ratio distortion, suppression will ultimately evolve, completely ($\lambda>1$) or partially ($\lambda=1$) restoring an equal sex ratio (Fig. 3aii & Fig. S6c & Fig. S6d). Consequently, we conclude that, with reasonable assumptions about the cost of suppression ($c_{sup}$), distorter suppression is the ultimate outcome of distorter evolution. For the sex ratio to be

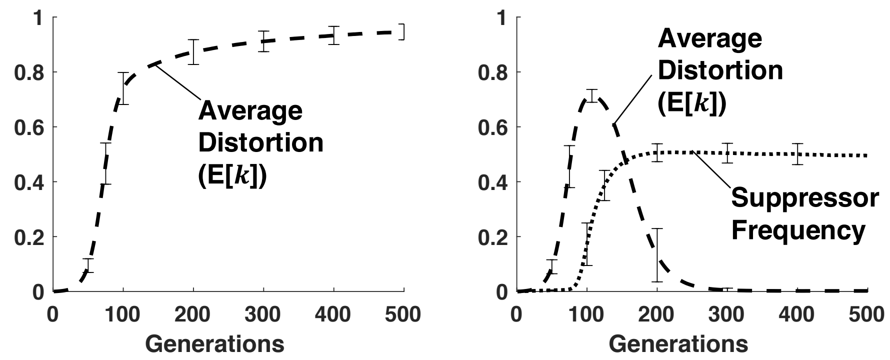


**Figure S7: Evolution of sex ratio distortion (dynamics).** This figure plots the results of the agent-based simulation model, in which X chromosome drive can mutate and take any value within 0-1. In (a), there is no suppressor allele (*sup*), and the population average level of X chromosome drive (E[$k$]) tends towards maximum distortion ($k$*=1). In (b), a suppressor of distortion (*sup*) is introduced from rarity. The population average level of X chromosome distortion (E[$k$]) increases alongside the suppressor (*sup*) frequency. Eventually, a threshold is passed, after which, distorting X chromosomes ($k_{x1}, k_{x2}>0$) are lost from the population. Sex ratio is restored to 0.5 at equilibrium. Double female mating ($\lambda$=2) and high suppression cost ($c_{\mathrm{sup}}$=0.3) were assumed in these simulations. The plots show average values over 100 runs, for $N$=100,000 individuals, with error bars plotting one standard deviation in each direction of the mean.

appreciably distorted (>60% females), suppression cost needs to exceed around $c_{sup}$=0.15 (λ=1) or $c_{sup}$=0.35 (λ>1) (Fig. S8).

By setting the frequencies of all genotypes bearing the suppressor (*sup*) to zero, we can use our recursions in Table S4 to find the equilibrium level of distortion in the absence of suppression. We exactly recover the equilibrium derived in Equation S4, which gives the ESS distortion ($k$*) in the absence of suppression (Fig. S6a).

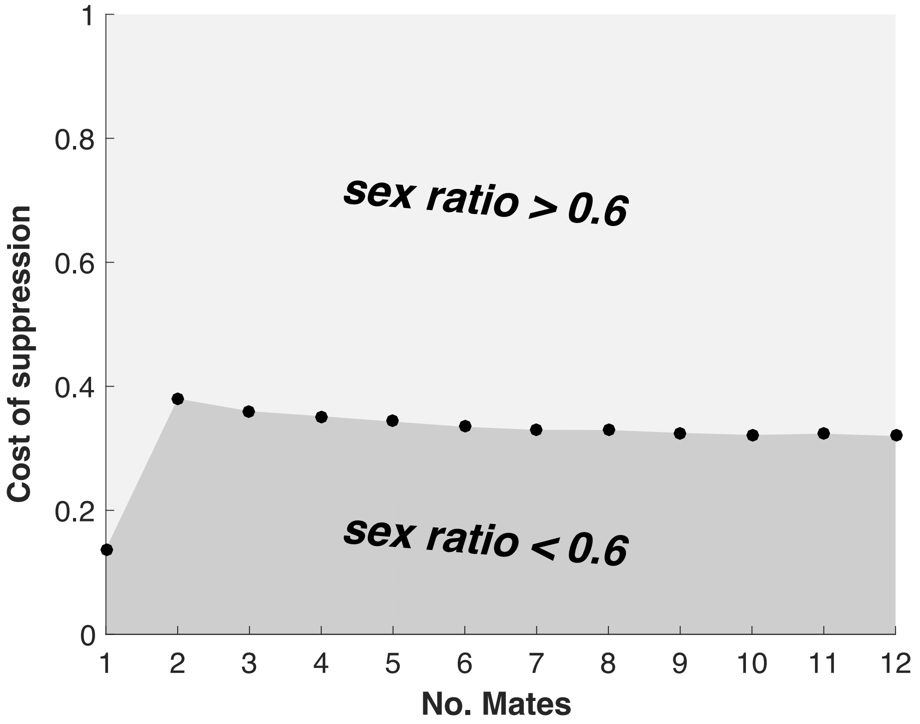


**Figure S8. Cost of suppression required for appreciable sex ratio distortion.** The equilibrium level of X chromosome distortion was obtained for different costs of suppression ($c_{sup}$). The sex ratio at this equilibrium level of X chromosome was recorded. The cost of suppression required for sex ratio distortion to be appreciably distorted (>60% females produced) is plotted (solid circles). Sex ratio is significantly distorted for the region above this curve.

#### Agent-based simulation

We construct an agent-based simulation to ask what level of sex ratio distortion evolves under strong selection, and when continuous variation is permitted at distorter and suppressor loci. We model a population of N=10,000 individuals and track evolution at an X chromosome distorter locus (L1) and an autosomal suppressor locus (L2). Individuals either have two alleles at the X chromosome locus, with strengths denoted by $k_{x1}$ and $k_{x2}$ (females), or one allele at the X chromosome locus, with strength denoted by $k_{x1}$ (males), and one Y chromosome.

Each allele at the distorter (L1) locus can take any continuous value between zero and one. Individuals have two alleles at the suppressor locus, with strengths denoted by $m_a$ and $m_b$ (diploid). At the suppressor (L2) locus, we consider both the case of: (i) discrete variation, in which suppressor strengths are either zero or one, and (ii) continuous variation, in which suppressor strengths can take any continuous value between zero and one. We assume that the strongest (highest value) suppressor allele within an individual is dominant. The ejaculate size of a given male is $1 - \frac{(1-\max(m_a, m_b))k_{x1}}{2}$, and his fertility is this value divided by the total sperm stored in the females it mates with. The total sperm store is the sum of ejaculates of this male and λ other males drawn at random from the population with replacement. The viability of males with an active distorter ($k_{x1}$>0) is given by 1-max($m_a$,$m_b$)$c_{sup}$. The viability of all other individuals is 1.

Each generation, there are N breeding pairs. Females are drawn at random with replacement to fill each female position in each breeding pair. Males are drawn from the population, with replacement, with probabilities given by their fertility. Breeding pairs then reproduce to produce one offspring, before dying (non-overlapping generations). Alleles at the suppressor locus (L2) are inherited in Mendelian fashion. Alleles at the distorter locus in males may drive, meaning the X chromosome is inherited, rather than the Y chromosome, with the probability (1+$k_{x1}$(1-max($m_a$,$m_b$)))/2. Offspring then compete for spots in the adult population, of which there are N. To fill each spot, offspring are drawn with replacement with likelihood that is proportional to their viability. Each generation, alleles at the distorter locus (L1), and for the case of continuous variation at the suppressor locus, alleles at the

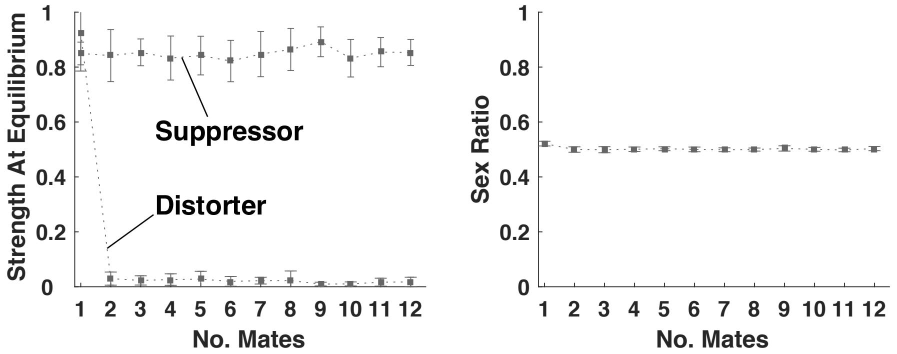


**Figure S9. Equilibrium distorter strength and sex ratio under continuous suppressor variation.** (a) Equilibrium suppressor strength (E[$m$*]) evolves to be high (~0.83). Maximally distorting (but largely suppressed) X chromosomes (E[$k$*]=1) evolve when females are singly mated ($\lambda$=1); otherwise, non-distorting X chromosomes (E[$k$*]=0) evolve. (b) Owing to the evolution of strong suppression, sex ratio is completely ($\lambda$>1) or partially ($\lambda$=1) recovered at equilibrium. Suppression cost in these simulations was $c_{sup}$=0.03. Error bars show one standard deviation from the mean over 10 trials.

suppressor locus (L2) have a 0.0005 chance of mutating to a new value, which is drawn from a normal distribution centred around the pre-mutation value, with variance 0.5, and truncated between 0 and 1 (strong selection). For the case of discrete variation at the suppressor locus, alleles at the suppressor locus (L2) have a 0.001 chance of mutating each generation between suppressor and non-suppressor states.

We iterate this lifecycle over 5,000 generations. We see that, when discrete variation is permitted at the suppressor locus (L2), the simulation quantitatively recovers the equilibrium level of distortion, and corresponding sex ratio, given by the game theoretic and population genetic models (Fig. S6). When continuous variation is permitted at the suppressor locus (L2), qualitatively equivalent results are obtained:

suppressor strength (E[*m*]) evolves to be high enough that sex ratio is fully (λ≥2) or partially (λ=1) recovered at equilibrium (Fig. S9).

### 5) Genomic Imprinting and Altruism

We consider an autosomal, maternally expressed selfish genetic element that may gain a propagative advantage by upregulating individual altruistic investment[46-51]. The genes that do not gain a propagative advantage from altruism upregulation comprise both paternally expressed and unimprinted genes. The conflict between maternally and paternally expressed genes, which can result in arms races and a 'tug of war' over organism phenotype, has been considered in previous theoretical work[52-55]. However, we focus on unimprinted suppressors, for simplicity, and because unimprinted genes comprise the larger group of genes (coreplicon), constituting the majority within the parliament of genes[56-58]. We focus our analyses on when a maternally expressed distorter and an unimprinted suppressor can spread. We first describe our modelling assumptions, then successively analyse the cases of unimprinted, and imprinted, altruism. The purpose of this model is to illustrate how selection will act on selfish imprinted genes and their suppressors.

**Modelling Assumptions**

We track a large population of diploid individuals. We consider a gene that induces an altruistic investment of some amount ($k>0$), at a fitness cost to the individual ($c(k)$), which is a monotonically increasing function of altruistic investment $\left(\frac{\partial c}{\partial k} \geq 0\right)$, and a benefit to the social partner ($b(k)>c(k)$), which is a function that is monotonically increasing with the level of altruistic investment $\left(\frac{\partial b}{\partial k} \geq 0\right)$ yet diminishing with the *cost* of altruistic investment $\left(\frac{d^2 b}{dc^2} \leq 0\right)$[59].

Each generation, male gametes (e.g. sperm) fuse at random with female gametes (e.g. eggs) to generate individuals (random mating). Individuals then pair up with other individuals who have matching maternally (egg-) inherited alleles at all loci; pairs are random with respect to identity at the paternally (sperm-) inherited allele at all loci ($R_m$=1; $R_p$=0). Individuals may then invest in altruism directed towards their partner, before producing gametes in proportion to their fitness (fertility), and dying (non-overlapping generations).

In nature, relatedness asymmetries within a generation may be generated by sex biased migration patterns[47], or as a consequence of greater variance in reproductive success in males[60,61]. They may alternatively be generated if kin recognition alleles are imprinted, which has been implicated in humans[62] and mice[63-65].

**Unimprinted Altruism**

We consider an unimprinted altruism gene, denoted by $y_A$, that, when homozygous, induces an altruistic investment of $k_A$ ($k_A>0$), and when heterozygous, induces an altruistic investment of $h*k_A$, where $h$ denotes the dominance. If we take $g$ and $g'$ as the population frequency of the altruism gene in two consecutive generations, then the population frequency of the altruism gene in the latter generation is:

$$\overline{w}g' = g^2\left(\frac{b(k_A)g(1-g)h}{g^2+g(1-g)} + \frac{b(k_A)g^2}{g^2+g(1-g)} - c(k_A) + 1\right) + \frac{1}{2}g(1-g)(\frac{g(1-g)b(k_A)h}{g^2+g(1-g)} + \frac{b(k_A)g^2}{g^2+g(1-g)} - c(k_A)h + 1) + \frac{1}{2}(1-g)g(\frac{b(k_A)(1-g)gh}{(1-g)^2+(1-g)g} - c(k_A)h + 1), \quad \text{(S5)}$$

where the mean fitness of individuals is given by:

$$\bar{w} = g(1-g)\left(\frac{g(1-g)b(k_A)h}{g^2+g(1-g)} + \frac{b(k_A)g^2}{g^2+g(1-g)} - c(k_A)h + 1\right) + g^2(\frac{g(1-g)b(k_A)h}{g^2+g(1-g)} + \frac{b(k_A)g^2}{g^2+g(1-g)} - c(k_A) + 1) + (1-g)g(\frac{(1-g)gb(k_A)h}{(1-g)^2+(1-g)g} - c(k_A)h + 1) + (1-g)^2(\frac{(1-g)gb(k_A)h}{(1-g)^2+(1-g)g} +1).$$

Each term relates to a different class of individual. For illustration, we derive the term relating to heterozygous individuals with a maternally derived $y_A$: $\frac{1}{2}g(1-g)\left(\frac{g(1-g)b(k_A)h}{g^2+g(1-g)} + \frac{b(k_A)g^2}{g^2+g(1-g)} - c(k_A)h + 1\right)$. (1) Find the frequency of individuals with this genotype: $g$*(1-$g$). (2) Multiply this by absolute fitness, which is 1 at baseline, with additively applied benefits weighted by the probability that the individual pairs with altruists, and additively applied costs applied if the individual is an altruist: $\frac{g(1-g)b(k_A)h}{g^2+g(1-g)} + \frac{b(k_A)g^2}{g^2+g(1-g)} - c(k_A)h + 1$. (3) Weight this by the proportion of $y_A$-bearing gametes produced by individuals: ½.

The altruism gene decreases in population frequency when $g' < g$, which requires the condition: ½b($k_A$)<c($k_A$). Given that genetic relatedness is $(R_f+R_m)/2$ = (0+1)/2 = ½, this condition corresponds to Hamilton's Rule[66-68]. This (½b($k_A$)<c($k_A$)) is also the condition for the invasion of a weaker altruism gene (lower $k_A$) against a stronger one, owing to diminishing returns on altruistic investment $\left(\frac{d^2b}{dc^2} < 0\right)$. Taken together, this implies that, when ½b($k_A$)<c($k_A$), the optimal altruism investment for unimprinted genes is zero, and increased altruistic investment is increasingly suboptimal.

**Selfish Gene Spread**

We consider an imprinted altruism gene that is only expressed when maternally inherited, denoted by $y_1$, and induces an altruistic investment of $k$ ($k$>0). If we take $p$ and $p'$ as the population frequency of the altruism gene in two consecutive generations, then the population frequency of the altruism gene in the latter generation is:

$$\bar{w}p' = p(1-p)(b(k)-c(k)+1)/2 + (1-p)p/2 + p^2(b(k)-c(k)+1), \qquad \text{(S6)}$$

where the mean fitness of individuals is given by:

$$\bar{w} = 1-p+(b(k)-c(k)+1)p.$$

Each term relates to a different class of individual. For illustration, we derive the term relating to heterozygous individuals with a paternally derived $y_1$: (1-$p$)$p$/2. (1) Find the population frequency of individuals with this genotype: (1-$p$)*$p$. (2) Weight by (absolute) individual fitness: 1. (3) Weight by the proportion of distorter-bearing gametes ($y_1$) produced: ½.

We ask when a rare imprinted altruism gene ($y_1$) can invade a population fixed for the non-distorter ($y_0$). We take Equation (S6), set $p'=p=p^*$, and solve to find two possible equilibria: $p^*=0$ (non-distorter fixation) and $p^*=1$ (imprinted gene fixation). The imprinted gene ($y_1$) can invade from rarity when the $p^*=0$ equilibrium is unstable, which occurs when the differential of *p*’ with respect to *p*, at $p^*=0$, is greater than one. The distorter invasion criterion is therefore $b(k)>c(k)$.

We now ask what frequency the imprinted altruism gene ($y_1$) will reach after invasion. The gene ($y_1$) can spread to fixation if the $p^*=1$ equilibrium is stable, which requires that the differential of *p*’ with respect to *p*, at $p^*=1$, is less than one. This requirement always holds true, demonstrating that there is no negative frequency dependence on the imprinted gene, and that it will always spread to fixation after its initial invasion.

Given that genetic relatedness is $R_m=1$, our condition for the spread of the imprinted altruism gene ($b(k)>c(k)$) corresponds to Hamilton’s Rule[66-68]. Combining with the result of the “Unimprinted altruism” model, altruistic investment (of $k_i=k_A=k$) is simultaneously favoured at maternally expressed genes and disfavoured at unimprinted genes, rendering the imprinted altruism gene a selfish trait distorter, when $\frac{1}{2}b(k_i)<c(k_i)<b(k_i)$.

**Spread of an autosomal suppressor**

We ask when an unimprinted suppressor (*sup*), competing against a non-suppressor (+), will invade from rarity. We can write recursions detailing the generational change in the frequencies of the four possible gametes, $y_0$/+, $y_0$/*sup*, $y_1$/+, $y_1$/*sup*, with the respective frequencies in the current generation denoted by $x_1$, $x_2$, $x_3$ and $x_4$, and the frequencies in the subsequent generation denoted by an appended dash (’):

$$\bar{w}x_1' = x_1x_1 + \frac{x_1x_2}{2} + \frac{x_1x_3}{2} + \frac{x_1x_4}{4} + \frac{x_2x_1}{2} + \frac{x_2x_3}{4} + \frac{1}{4}x_4x_1(1 - c_{\text{sup}}) + \frac{1}{4}x_3x_2(1 - c_{\text{sup}})(1$$
$$+b(k)(x_1 + x_3)) + \frac{1}{2}x_3x_1(1 - c(k) + b(k)(x_1 + x_3))$$

$$\bar{w}x_2' = \frac{x_1}{4} + \frac{x_1x_2}{2} + \frac{x_1x_4}{4} + \frac{x_2x_1}{2} + x_2x_2 + \frac{x_2x_3}{4} + \frac{x_2x_4}{2} + \frac{1}{4}x_4x_1(1 - c_{\text{sup}}) + \frac{1}{2}x_4x_2(1$$
$$-c_{\text{sup}}) + \frac{1}{4}x_3x_2(1 - c_{\text{sup}})(1 + b(k)(x_1 + x_3))$$

$$\bar{w}x_3' = \frac{x_1x_3}{2} + \frac{x_1x_4}{4} + \frac{x_2x_3}{4} + \frac{1}{4}x_4x_1\left(1 - c_{\text{sup}}\right) + \frac{1}{2}x_4x_3\left(1 - c_{\text{sup}}\right) + \frac{1}{4}x_3x_2\left(1 - c_{\text{sup}}\right)$$
$$(1 + b(k)(x_1 + x_3)) + \frac{1}{2}x_3x_4(1 - c_{\text{sup}})(1 + b(k)(x_1 + x_3)) + \frac{1}{2}x_3x_1(1 -$$
$$c(k) + b(k)(x_1 + x_3)) + x_3x_3(1 - c(k) + b(k)(x_1 + x_3))$$

$$\bar{w}x_4' = \frac{x_1x_4}{4} + \frac{x_2x_3}{4} + \frac{x_2x_4}{2} + \frac{1}{4}x_4x_1\left(1 - c_{\text{sup}}\right) + \frac{1}{2}x_4x_2\left(1 - c_{\text{sup}}\right) + \frac{1}{2}x_4x_3\left(1 - c_{\text{sup}}\right)$$
$$+x_4x_4(1 - c_{\text{sup}}) + \frac{1}{4}x_3x_2(1 - c_{\text{sup}})(1 + b(k)(x_1 + x_3)) + \frac{1}{2}x_3x_4(1 - c_{\text{sup}})(1$$
$$+b(k)(x_1 + x_3)) \qquad \text{(S7)}$$

$\bar{w}$ is the average fitness of individuals in the current generation, and equals the sum of the equations' right-hand sides. Each term in each equation relates to a different class of individual. For illustration, we derive the term corresponding to the contribution of $y_0$/+ gametes to the next generation, by individuals with a maternally inherited $y_1$/+ gamete and a paternally inherited $y_0$/+ gamete; this is the $\frac{1}{2}x_3x_1(1 - c(k) + b(k)(x_1 + x_3))$ term in the $\bar{w}x_1'$ recursion. (1) Find the population frequency of individuals with this genotype: $x_3x_1$. (2) Weight by (absolute) individual fitness: $1 - c(k) + b(k)(x_1 + x_3)$. (3) Weight by the proportion of distorter-bearing gametes ($y_1$) produced: ½.

We derive the Jacobian stability matrix for the equilibrium in which the distorter ($y_1$) and non-suppressor (+) are at fixation ($x_1$*=0, $x_2$*=0, $x_3$*=1, $x_4$*=0). The suppressor can invade when the equilibrium is unstable, which occurs when the leading eigenvalue is greater than one. The leading eigenvalue is $\frac{(b(k)+2)(1-c_{sup})}{2(b(k)-c(k)+1)}$, meaning the suppressor invasion criterion is given by:

$$c_{sup}(1+b(k)/2)<c(k)-b(k)/2. \quad \text{(S8)}$$

Therefore, the suppressor invades from rarity above a threshold level of distortion, $k$, when, from the perspective of an unimprinted locus, the number of relatives that die as a result of trait distortion ($c(k)$-$b(k)/2$), exceeds the number of relatives that die as a result of distorter suppression ($c_{sup}(1+b(k)/2)$).

**Consequences of suppressor spread for organism phenotype**

We ask what frequency the distorter ($y_1$) and suppressor (*sup*) will reach after initial suppressor (*sup*) invasion. We assume that the suppressor is introduced from rarity when the distorter has reached the population frequency given by $f$ (x1→$f$, x3→1-$f$, {x2,x4}→0). We numerically iterate Equations (S7), over successive generations, until equilibrium has been reached. At equilibrium, for all parameter combinations ($f,t,c_{sup},c_{drive}$), the suppressor reaches an internal equilibrium and the distorter is lost from the population ($x_1^*+x_2^*=1$, $x_3^*=0$, $x_4^*=0$). This equilibrium arises because distorter-presence gives the suppressor (*sup*) a selective advantage, leading to high suppressor frequency, which in turn reverses the selective advantage of the distorter ($y_1$), leading to distorter loss and suppressor equilibration (Fig. 3bi).

**Invasion of a mutant distorter**

We ask when a mutant distorter (*y*$_2$) of strength $\hat{k}$ will invade against a resident distorter (*y*$_1$) that is unsuppressed and at fixation ($\hat{k}\neq k$). We write recursions detailing the generational frequency changes in the six possible gametes, $y_0/+$, $y_0/sup$, $y_1/+$, $y_1/sup$, $y_2/+$, $y_2/sup$, with current generation frequencies denoted respectively by $x_1$, $x_2$, $x_3$, $x_4$, $x_5$, $x_6$, and next generation frequencies denoted with an appended dash ('):

$$\bar{w}x_1' = x_1x_1 + \frac{x_1x_2}{2} + \frac{x_1x_3}{2} + \frac{x_1x_4}{4} + \frac{x_1x_5}{2} + \frac{x_1x_6}{4} + \frac{x_2x_1}{2} + \frac{x_2x_3}{4} + \frac{x_2x_5}{4} + \frac{1}{4}x_4x_1\left(1 - c_{sup}\right)$$

$$+\frac{1}{4}x_6x_1(1 - c_{sup}) + \frac{1}{4}x_3x_2(1 - c_{sup})(1 + b(k)(x_1 + x_3 + x_5)) + \frac{1}{2}x_3x_1(1 - c(k)$$

$$+b(k)(x_1 + x_3 + x_5)) + \frac{1}{4}x_5x_2(1 - c_{sup})(1 + b(\hat{k})(x_1 + x_3 + x_5)) + \frac{1}{2}x_5x_1(1 -$$

$$c(\hat{k}) + b(\hat{k})(x_1 + x_3 + x_5))$$

$$\bar{w}x_2' = \frac{x_1x_2}{2} + \frac{x_1x_4}{4} + \frac{x_1x_6}{4} + \frac{x_2x_1}{2} + x_2x_2 + \frac{x_2x_3}{4} + \frac{x_2x_4}{2} + \frac{x_2x_5}{4} + \frac{x_2x_6}{2} + \frac{1}{4}x_4x_1\left(1 - c_{sup}\right)$$

$$+\frac{1}{2}x_4x_2(1 - c_{sup}) + \frac{1}{4}x_6x_1(1 - c_{sup}) + \frac{1}{2}x_6x_2(1 - c_{sup}) + \frac{1}{4}x_3x_2(1 - c_{sup})(1$$

$$+b(k)(x_1 + x_3 + x_5)) + \frac{1}{4}x_5x_2(1 - c_{sup})(1 + b(\hat{k})(x_1 + x_3 + x_5))$$

$$\bar{w}x_3' = \frac{x_1x_3}{2} + \frac{x_1x_4}{4} + \frac{x_2x_3}{4} + \frac{1}{4}x_4x_1\left(1 - c_{sup}\right) + \frac{1}{2}x_4x_3\left(1 - c_{sup}\right) + \frac{1}{4}x_4x_5\left(1 - c_{sup}\right)$$

$$+\frac{1}{4}x_6x_3(1 - c_{sup}) + \frac{1}{4}x_3x_2(1 - c_{sup})(1 + b(k)(x_1 + x_3 + x_5)) + \frac{1}{2}x_3x_4(1 -$$

$$c_{sup})(1 + b(k)(x_1 + x_3 + x_5)) + \frac{1}{4}x_3x_6(1 - c_{sup})(1 + b(k)(x_1 + x_3 + x_5)) +$$

$$\frac{1}{2}x_3x_1(1 - c(k) + b(k)(x_1 + x_3 + x_5)) + x_3x_3(1 - c(k) + b(k)(x_1 + x_3 +$$

$$x_5)) + \frac{1}{2}x_3x_5(1 - c(k) + b(k)(x_1 + x_3 + x_5)) + \frac{1}{4}x_5x_4(1 - c_{sup})(1 +$$

$$b(\hat{k})(x_1 + x_3 + x_5)) + \frac{1}{2}x_5x_3(1 - c(\hat{k}) + b(\hat{k})(x_1 + x_3 + x_5))$$

$$\bar{w}x_4' = \frac{x_1x_4}{4} + \frac{x_2x_3}{4} + \frac{x_2x_4}{2} + \frac{1}{4}x_4x_1\left(1 - c_{sup}\right) + \frac{1}{2}x_4x_2\left(1 - c_{sup}\right) + \frac{1}{2}x_4x_3\left(1 - c_{sup}\right)$$

$$+x_4x_4\left(1 - c_{sup}\right) + \frac{1}{4}x_4x_5\left(1 - c_{sup}\right) + \frac{1}{2}x_4x_6\left(1 - c_{sup}\right) + \frac{1}{4}x_6x_3\left(1 - c_{sup}\right)$$

$$+\frac{1}{2}x_6x_4(1-c_{sup})+\frac{1}{4}x_3x_2(1-c_{sup})(1+b(k)(x_1+x_3+x_5))+\frac{1}{2}x_3x_4(1-c_{sup})(1+b(k)(x_1+x_3+x_5))+\frac{1}{4}x_3x_6(1-c_{sup})(1+b(k)(x_1+x_3+x_5))+\frac{1}{4}x_5x_4(1-c_{sup})(1+b(\hat{k})(x_1+x_3+x_5))$$

$$\bar{w}x_5{}'=\frac{x_1x_5}{2}+\frac{x_1x_6}{4}+\frac{x_2x_5}{4}+\frac{1}{4}x_4x_5\left(1-c_{sup}\right)+\frac{1}{4}x_6x_1\left(1-c_{sup}\right)+\frac{1}{4}x_6x_3\left(1-c_{sup}\right)+\frac{1}{2}x_6x_5(1-c_{sup})+\frac{1}{4}x_3x_6(1-c_{sup})(1+b(k)(x_1+x_3+x_5))+\frac{1}{2}x_3x_5(1-c(k)+b(k)(x_1+x_3+x_5))+\frac{1}{4}x_5x_2(1-c_{sup})(1+b(\hat{k})(x_1+x_3+x_5))+\frac{1}{4}x_5x_4(1-c_{sup})(1+b(\hat{k})(x_1+x_3+x_5))+\frac{1}{2}x_5x_6(1-c_{sup})(1+b(\hat{k})(x_1+x_3+x_5))+\frac{1}{2}x_5x_1(1-c(\hat{k})+b(\hat{k})(x_1+x_3+x_5))+\frac{1}{2}x_5x_3(1-c(\hat{k})+b(\hat{k})(x_1+x_3+x_5))+x_5x_5(1-c(\hat{k})+b(\hat{k})(x_1+x_3+x_5))$$

$$\bar{w}x_6{}'=\frac{x_1x_6}{4}+\frac{x_2x_5}{4}+\frac{x_2x_6}{2}+\frac{1}{4}x_4x_5\left(1-c_{sup}\right)+\frac{1}{2}x_4x_6\left(1-c_{sup}\right)+\frac{1}{4}x_6x_1\left(1-c_{sup}\right)+\frac{1}{2}x_6x_2\left(1-c_{sup}\right)+\frac{1}{4}x_6x_3\left(1-c_{sup}\right)+\frac{1}{2}x_6x_4\left(1-c_{sup}\right)+\frac{1}{2}x_6x_5\left(1-c_{sup}\right)+x_6x_6(1-c_{sup})+\frac{1}{4}x_3x_6(1-c_{sup})(1+b(k)(x_1+x_3+x_5))+\frac{1}{4}x_5x_2(1-c_{sup})(1+b(\hat{k})(x_1+x_3+x_5))+\frac{1}{4}x_5x_4(1-c_{sup})(1+b(\hat{k})(x_1+x_3+x_5))+\frac{1}{2}x_5x_6(1-c_{sup})(1+b(\hat{k})(x_1+x_3+x_5)). \qquad \text{(S9)}$$

$\bar{w}$ is the average fitness of individuals in the current generation, and equals the sum of the right-hand side of the system of equations. The mutant distorter can invade when the equilibrium given by $x_1^*=0$, $x_2^*=0$, $x_3^*=1$, $x_4^*=0$, $x_5^*=0$, $x_6^*=0$ is unstable, which occurs when the leading eigenvalue of the Jacobian stability matrix for this equilibrium is greater than one. Testing for stability in this way, we find that the mutant distorter invades from rarity when $\Delta b>\Delta c$, where $\Delta b=b(\hat{k})-b(k)$, $\Delta c=c(\hat{k})-c(k)$.

The implication is that mutant distorters will invade if they approach a 'target' strength ($k_{target}$) at which:

$$\frac{\partial b}{\partial k} = \frac{\partial c}{\partial k}. \qquad \text{(S10)}$$

In the absence of suppression, this target is the equilibrium level of distortion ($k^*=k_{target}$).

**Equilibrium distorter and suppressor frequencies (long term evolution)**

We ask what equilibrium state will arise after the invasion of a mutant distorter. We assume that the mutant distorter ($y_2$) is introduced from rarity when the resident distorter ($y_1$) has reached the population frequency given by $q$. We numerically iterate Equations (S9), over successive generations, until equilibrium has been reached. At equilibrium, for all parameter combinations ($q,t(k),t(\hat{k}),c_{sup},c(k),c(\hat{k})$), the resident distorter ($y_1$) is lost from the population ($x_3,x_4=0$), with either the mutant distorter ($y_2$) and non-suppressor (+) at fixation ($x_5^*=1$), or the non-distorter at fixation alongside the suppressor at an internal equilibrium ($x_1^*+x_2^*=1$). The latter scenario arises if the mutant distorter triggers suppressor invasion ($c_{sup}(1+b(\hat{k})/2)<c(\hat{k})-b(\hat{k})/2$). This equilibrium arises because mutant distorter-presence gives the suppressor (*sup*) a selective advantage, leading to high suppressor frequency, which in turn reverses the selective advantage of distortion, leading to distorter ($y_1,y_2$) loss and suppressor equilibration.

Given that mutant distorters will invade if they approach a 'target' strength ($k_{target}$), if the individual level cost associated with this target level of distortion ($c(k_{\mathrm{target}})$) is sufficiently high relative to the cost of suppression ($c_{sup}$), so that the following condition is satisfied, the equilibrium level of distortion will be $k^*=0$: $c_{sup}(1+b(k_{\mathrm{target}})/2)<c(k_{\mathrm{target}})-b(k_{\mathrm{target}})/2$. If this condition is not satisfied the equilibrium level of distortion will be $k^*=k_{target}$ (Fig. 3bii).

#### 6) Horizontal Gene Transfer and Public Goods

We model the evolution of investment in a public good in a large, clonally reproducing population. We assume a public good that costs *c* to produce, and provides a benefit *b* to the group. We assume a well-mixed population, meaning genetic relatedness at vertically inherited genes is zero ($R_{vertical}$=0), and so indirect fitness benefits cannot favour public good production at the individual level ($R_{vertical}b=0<c$)[66-70]. There are also direct fitness benefits of public good production, which arise because producers of public goods receive a fraction of the benefit (*b*) they confer on the group, but we assume that the population is sufficiently large and well mixed that direct fitness benefits cannot favour public good production at the individual level. This means that public good production is disfavoured at the individual level.

We consider a selfish genetic element that resides on a mobile locus (horizontal & vertical transmission) and may gain a propagative advantage by upregulating individual public goods investment[71-75]. The genes that do not gain a propagative advantage from increased public goods production comprise the non-mobile loci (vertical transmission). Non-mobile loci comprise most of the genome, and so constitute the majority within the parliament of genes. We focus our analyses on when a mobile distorter and a non-mobile suppressor can spread. The purpose of this model is to illustrate how selection will act on selfish mobile genes and their suppressors.

**Model assumptions**

We consider a public goods gene ($y_1$) that competes against a non-distorter ($y_0$) at a mobile locus. The distorter ($y_1$) increases public goods investment by some amount ($k$), at a fitness cost to the individual ($c(k)$) and benefit shared within the group ($b(k)>c(k)$) that are both monotonically increasing functions of investment $\left(\frac{\partial\{b,c\}}{\partial k} \geq 0\right)$.

We assume the following lifecycle. Individuals in a large, effectively infinite, population randomly aggregate into smaller social groups (*patches*). Individuals then randomly pair up within their patch, and horizontal gene transfer occurs, with certainty, within pairs that are genetically dissimilar at the mobile locus[76,77]. Alternative assumptions about the probability of horizontal gene transfer do not change our qualitative results (Scott, unpublished). Only one allele at the mobile locus is transferrable in each patch, and each allele at the mobile locus is transferrable in an equal proportion of patches. We denote those patches in which the non-distorter ($y_0$) is transferred as "type 1" patches, and those patches in which the distorter ($y_1$) is transferred as "type 2" patches. Individuals may then produce public goods, which are shared within patches, before the population re-merges, and individuals reproduce in proportion to their fitness before dying (non-overlapping generations), with progeny inheriting all alleles from their parent (perfect inheritance).

**Selfish Gene Spread**

We respectively take $\bar{p}$ and $\bar{p}''$ as the population frequency of the distorter ($y_1$) at the start of two consecutive generations, and $p'_j$ as the average frequency of the distorter ($y_1$) in patches of type $j$ after horizontal gene transfer ($j \in \{1,2\}$), with $p'_{j=1} = \bar{p}$

+ $\bar{p}$(1-$\bar{p}$) and $p'_{j=2}$ = $\bar{p}$ – $\bar{p}$(1-$\bar{p}$). The population frequency of the distorter in the latter generation ($\bar{p}''$) is:

$$\bar{p}'' = \frac{p'_{j=1}\left(1+p'_{j=1}b(k)-c(k)\right)+p'_{j=2}\left(1+p'_{j=2}b(k)-c(k)\right)}{2+(b(k)-c(k))\left(p'_{j=1}+p'_{j=2}\right)}, \tag{S11}$$

where the denominator denotes average individual fitness. Stable equilibria occur for $\bar{p}$=$\bar{p}''$=$p^*$ and $\frac{\partial \bar{p}''}{\partial \bar{p}}\Big|_{\bar{p}=p^*} < 1$, which occurs when $p^* = \left\{0, \left(1+\sqrt{1-\frac{4c(k)}{b(k)}}\right)/2\right\}$. Unstable equilibria occur for $\bar{p}$=$\bar{p}''$=$\bar{p}^*$ and $\frac{\partial \bar{p}''}{\partial \bar{p}}\Big|_{\bar{p}=p^*} > 1$, which occurs when $p^* = \left\{\left(1-\sqrt{1-\frac{4c(k)}{b(k)}}\right)/2, 1\right\}$. Therefore, the distorter ($y_1$) exhibits positive and negative frequency dependence, meaning it can only invade if introduced at high enough frequency $\left(\bar{p} > \left(1-\sqrt{1-\frac{4c(k)}{b(k)}}\right)/2\right)$, reaching a polymorphism below fixation $\left(p^* = \left(1+\sqrt{1-\frac{4c(k)}{b(k)}}\right)/2\right)$ (Fig. S10). Frequency dependence arises because, when there is low genetic diversity at the mobile locus ($p$→0/1), there is less generational horizontal gene transfer, and correspondingly lower patch relatedness, which dissipates the distorter's selective advantage[71]. A distorter ($y_1$) is more likely to invade and reach a high population frequency if it produces a public good associated with a large benefit to cost ratio (*b*/*c*)[71-73,78-80].

**Spread of a suppressor and consequences for the organism**

We consider a suppressor allele (*sup*) that competes against a non-suppressor (+) at a non-mobile locus. Suppressors of mobile elements are widespread and may

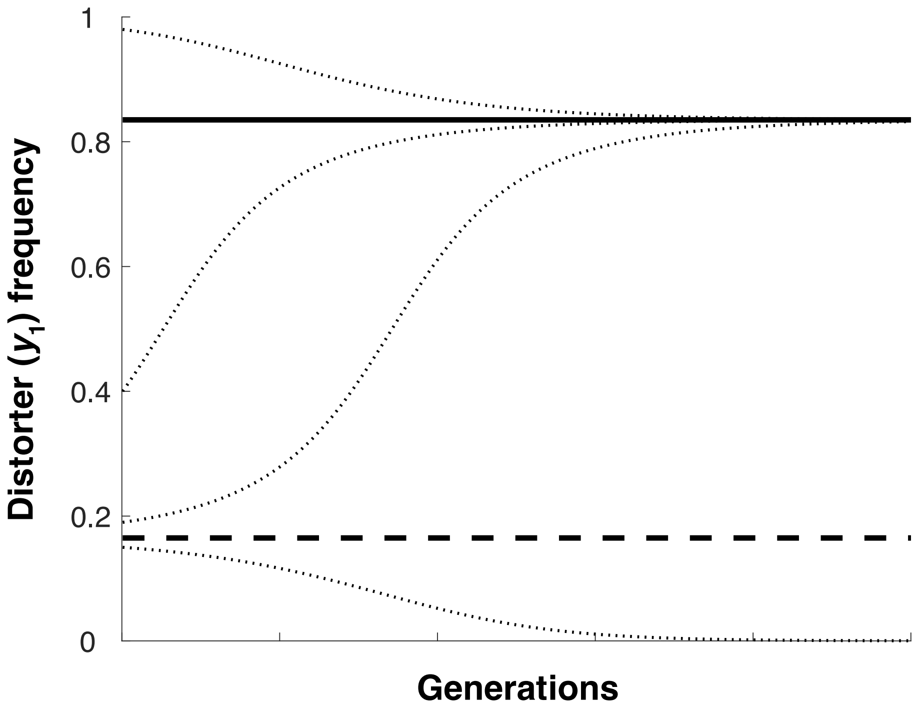


**Figure S10. Spread of the distorter ($y_1$) in the absence of suppression.** The distorter ($y_1$) is introduced at some frequency, and equilibrates over successive generations (dotted lines indicate trajectories corresponding to different initial distorter frequencies). The solid and dashed lines are, respectively, the stable $\left(p^* = \left(1 + \sqrt{1 - \frac{4c(k)}{b(k)}}\right)/2\right)$ and unstable $\left(p^* = \left(1 - \sqrt{1 - \frac{4c(k)}{b(k)}}\right)/2\right)$ equilibria. If introduced at a frequency greater than the unstable internal equilibrium, distorter ($y_1$) frequency reaches the stable internal equilibrium, without going to fixation (negative frequency dependence). If introduced at a frequency lower than the unstable internal equilibrium, distorter ($y_1$) frequency goes to zero (positive frequency dependence).

silence elements before they are translated, through gene methylation and RNAi[81]. We respectively take $\bar{x}_i$ and $\bar{x}_i''$ as the population genotype frequencies at the start of two consecutive generations, with the subscript $i \in \{1,2,3,4\}$ denoting the respective genotypes: $\{y_0/+, y_0/sup, y_1/+, y_1/sup\}$. We take $x_{ij}'$ as the average frequency of genotype $i$ in patches of type $j$ after horizontal gene transfer ($j \in \{1,2\}$), with $x_{11}' = \bar{x}_{i=1} + (\bar{x}_{i=1} + \bar{x}_{i=2})\bar{x}_{i=3}$, $x_{21}' = \bar{x}_{i=2} + (\bar{x}_{i=1} + \bar{x}_{i=2})\bar{x}_{i=4}$, $x_{31}' = \bar{x}_{i=3} - (\bar{x}_{i=1} + \bar{x}_{i=2})\bar{x}_{i=3}$, $x_{41}' = \bar{x}_{i=4} - (\bar{x}_{i=1} + \bar{x}_{i=2})\bar{x}_{i=4}$, $x_{12}' = \bar{x}_{i=1} - (\bar{x}_{i=3} + \bar{x}_{i=4})\bar{x}_{i=1}$, $x_{22}' =$

$\bar{x}_{i=2} - (\bar{x}_{i=3} + \bar{x}_{i=4})\bar{x}_{i=2}$, $x_{32}{}' = \bar{x}_{i=3} + (\bar{x}_{i=3} + \bar{x}_{i=4})\bar{x}_{i=1}$, $x_{42}{}' = \bar{x}_{i=4} + (\bar{x}_{i=3} + \bar{x}_{i=4})\bar{x}_{i=2}$. The population genotype frequencies in the latter generation ($\bar{x}_i{}'$) are:

$$W\bar{x}_1{}'' = \sum_{j=1}^{j=2}(x_{1j}{}'(1 + x_{3j}{}'b(k))),$$

$$W\bar{x}_2{}'' = \sum_{j=1}^{j=2}(x_{2j}{}'(1 + x_{3j}{}'b(k))),$$

$$W\bar{x}_3{}'' = \sum_{j=1}^{j=2}(x_{3j}{}'(1 + x_{3j}{}'b(k) - c(k))),$$

$$W\bar{x}_4{}'' = \sum_{j=1}^{j=2}(x_{4j}{}'(1 + x_{3j}{}'b(k))(1 - c_{sup}))), \quad \text{(S12)}$$

where $W$ is average individual fitness, equal to the sum of the right-hand sides of the system of equations.

We numerically iterated these recursions, for a range of parameter values ($b$,$c$,$c_{sup}$), and for different initial frequencies of the distorter ($y_1$) to find the distorter ($y_1$) and suppressor (*sup*) frequencies at equilibrium, and the resulting average trait distortion ($x_3\, k$). We found that, when distortion is weak (low $k$), suppressors are not favoured, but the distorter has relatively little impact at the individual level. For example, when the cost of suppression is $c_{sup}$=0.05, and the cost and benefit of public goods production are $c_{HGT}$=$k$ (linear cost) and $b_{HGT}$=8$k^{0.9}$ (relatively large, decelerating benefit), unsuppressed distorters cannot upregulate public goods by more than $k$=$c_{HGT}$=0.08 (Fig. 3ci).

We found that the suppressor invades from rarity, in response to a distorter at equilibrium $\left(x_3^* = \left(1 + \sqrt{1 - \frac{4c(k)}{b(k)}}\right)/2\right)$, above a threshold level of distortion. If the suppressor invades, it increases in frequency until the distorter's ($y_1$) selective advantage is reversed and the distorter is lost from the population; the suppressor (*sup*) then equilibrates (Fig. 3ci). A distorter ($y_1$) is more likely to evade suppression if it produces a public good associated with a large benefit to cost ratio ($b(k)/c(k)$) and if there is a high cost of suppression ($c_{sup}$)[82].

**Evolution of trait distortion**

We ask when a mutant distorter ($y_2$) of strength ($\hat{k}$) will invade against a resident distorter ($y_1$) that is unsuppressed and at equilibrium ($\hat{k} \neq k$). We denote those patches in which the mutant distorter ($y_2$) is transferred as "type 3" patches. We use the subscript $i \in \{1,2,3,4,5,6\}$ to denote the respective genotypes $\{y_0/+, y_0/sup, y_1/+, y_1/sup, y_2/+, y_2/sup\}$, and $j \in \{1,2,3\}$ to denote patch type. Average genotype frequencies in each patch type after horizontal gene transfer ($x_{ij}'$) are given by: $x_{11}' = \bar{x}_{i=1} + (\bar{x}_{i=1} + \bar{x}_{i=2})(\bar{x}_{i=3} + \bar{x}_{i=5})$, $x_{21}' = \bar{x}_{i=2} + (\bar{x}_{i=1} + \bar{x}_{i=2})(\bar{x}_{i=4} + \bar{x}_{i=6})$, $x_{31}' = \bar{x}_{i=3} - (\bar{x}_{i=1} + \bar{x}_{i=2})\bar{x}_{i=3}$, $x_{41}' = \bar{x}_{i=4} - (\bar{x}_{i=1} + \bar{x}_{i=2})\bar{x}_{i=4}$, $x_{51}' = \bar{x}_{i=5} - (\bar{x}_{i=1} + \bar{x}_{i=2})\bar{x}_{i=5}$, $x_{61}' = \bar{x}_{i=6} - (\bar{x}_{i=1} + \bar{x}_{i=2})\bar{x}_{i=6}$, $x_{12}' = \bar{x}_{i=1} - (\bar{x}_{i=3} + \bar{x}_{i=4})\bar{x}_{i=1}$, $x_{22}' = \bar{x}_{i=2} - (\bar{x}_{i=3} + \bar{x}_{i=4})\bar{x}_{i=2}$, $x_{32}' = \bar{x}_{i=3} + (\bar{x}_{i=3} + \bar{x}_{i=4})(\bar{x}_{i=1} + \bar{x}_{i=5})$, $x_{42}' = \bar{x}_{i=4} + (\bar{x}_{i=3} + \bar{x}_{i=4})(\bar{x}_{i=2} + \bar{x}_{i=6})$, $x_{52}' = \bar{x}_{i=5} - (\bar{x}_{i=3} + \bar{x}_{i=4})\bar{x}_{i=5}$, $x_{62}' = \bar{x}_{i=6} - (\bar{x}_{i=3} + \bar{x}_{i=4})\bar{x}_{i=6}$, $x_{13}' = \bar{x}_{i=3} - (\bar{x}_{i=5} + \bar{x}_{i=6})\bar{x}_{i=1}$, $x_{23}' = \bar{x}_{i=2} - (\bar{x}_{i=5} + \bar{x}_{i=6})\bar{x}_{i=2}$, $x_{33}' = \bar{x}_{i=3} - (\bar{x}_{i=5} + \bar{x}_{i=6})\bar{x}_{i=3}$, $x_{43}' = \bar{x}_{i=4} - (\bar{x}_{i=5} + \bar{x}_{i=6})\bar{x}_{i=4}$, $x_{51}' = \bar{x}_{i=5} + (\bar{x}_{i=5} + \bar{x}_{i=6})(\bar{x}_{i=1} + \bar{x}_{i=3})$, $x_{61}' = \bar{x}_{i=6} + (\bar{x}_{i=5} + \bar{x}_{i=6})(\bar{x}_{i=2} + \bar{x}_{i=4})$. We write recursions detailing the generational genotype frequency changes:

$$W\bar{x}_1'' = \sum_{j=1}^{j=3}(x_{1j}'(1 + x_{3j}'b(k) + x_{5j}'b(\hat{k}))),$$

$$W\bar{x}_2'' = \sum_{j=1}^{j=3}(x_{2j}'(1 + x_{3j}'b(k) + x_{5j}'b(\hat{k}))),$$

$$W\bar{x}_3'' = \sum_{j=1}^{j=3}(x_{3j}'(1 + x_{3j}'b(k) + x_{5j}'b(\hat{k}) - c(k))),$$

$$W\bar{x}_4'' = \sum_{j=1}^{j=3}(x_{4j}'(1 + x_{3j}'b(k) + x_{5j}'b(\hat{k}))(1 - c_{sup}))),$$

$$W\bar{x}_5'' = \sum_{j=1}^{j=3}(x_{5j}'(1 + x_{3j}'b(k) + x_{5j}'b(\hat{k}) - c(\hat{k}))),$$

$$W\bar{x}_6'' = \sum_{j=1}^{j=3}(x_{6j}'(1 + x_{3j}'b(k) + x_{5j}'b(\hat{k}))(1 - c_{sup}))), \qquad \text{(S13)}$$

where *W* is average individual fitness, equal to the sum of the right-hand sides of the system of equations.

We assume that distortion ($k$) is initially low, and introduce successive mutant distorters ($y_2$), each deviating only slightly from the distorters from which they are derived, until one fails to displace the resident distorter. The strength of the non-invadable allele gives the equilibrium level of distortion under $\delta$-weak selection[45]. We find that, if the rate of decrease in marginal cooperative benefits $\left(-\frac{d^2b}{dk^2}\right)$ is high relative to the rate of increase in marginal cooperative costs $\left(\frac{d^2c}{dk^2}\right)$, distortion ($k^*$) evolves to be low, and the suppressor (*sup*) may not invade. Otherwise, stronger distorters ($y_2$) successively invade, bringing distortion above the threshold level at which the suppressor (*sup*) spreads, with the end result that distorters are suppressed and lost from the population, with no trait distortion at equilibrium ($k^*$=0) (Fig. 3cii).

We lack empirical data that would allow us to test our model of mobile public goods genes. Genes associated with extracellular traits, which could represent cooperative public goods, appear to be overrepresented on mobile elements[80]. However, this may be nothing to do with cooperation *per se* – genes involved with adaptation to new environments might be more likely to be horizontally acquired, and extracellular traits might be especially important in adaptation to new environments[73-76,83].

### 7) Additional Discussion

**Suppressors**

We assumed facultative suppressors, where the cost of suppression is only incurred whilst the distorter is present. If the cost of the suppressor arises irrespective of whether a distorter is present (obligate), this could make it harder for suppressors to invade, but there are clear mechanisms by which facultative suppression can evolve[29,41,84,85], and facultative suppressors would be able to outcompete obligate ones.

**Dynamics and Mutation**

In nature, at any one point in time, the extent to which individual-level traits will be distorted by selfish genetic elements will depend on the rate at which trait-distorting selfish genetic elements arise by mutation, relative to the rate at which their suppressors arise by mutation. Given that the coreplicons that can spawn trait distorters are vastly smaller than the coreplicons that can spawn suppressors, we would expect, in general, that a given individual would have very few unsuppressed selfish genetic elements at any given point in time[2]. In support of this, whole genome sequencing of Arabidopsis lineages has revealed that novel segregation distorters evolve rapidly (~1 per 244,000 years), but distorters are only revealed in hybrid crosses, implying that suppressors have evolved rapidly in response[86,87].

**Individual Fitness Maximisation and the Parliament of Genes**

Our analysis suggests solutions for several problems that have been levelled at individual level fitness maximisation and the parliament of genes[88-93]. Ridley[94] argued that adaptation may be limited if large portions of the genome are wasted by embroilment in “subversion and counter subversion”, and that segregation at suppressor loci might expose previously suppressed distorters. Our analysis shows that distorters will generally be purged once their selective advantage is reversed by suppression, freeing up genomic space, and minimising their risk of becoming cross-generationally re-expressed. Crow[44] and others have pointed out that the cost associated with some selfish genetic elements stems from linked deleterious genes, which is not recovered via suppression of the selfish genetic element, in which case suppressors will not spread in response to drivers that reach fixation[95]. However, such drivers do not systematically bias trait values, and so pose no problem to individual level fitness maximisation. We have shown that when traits are distorted, selection for suppressors is retained after driver fixation.
